# Fast segmentation with the NextBrain histological atlas

**DOI:** 10.1101/2025.09.22.673638

**Authors:** Oula Puonti, Jackson Nolan, Robert Dicamillo, Yael Balbastre, Adria Casamitjana, Matteo Mancini, Eleanor Robinson, Loic Peter, Roberto Annunziata, Juri Althonayan, Shauna Crampsie, Emily Blackburn, Benjamin Billot, Alessia Atzeni, Peter Schmidt, James Hughes, Jean Augustinack, Brian Edlow, Lilla Zöllei, David L Thomas, Dorit Kliemann, Martina Bocchetta, Catherine Strand, Janice Holton, Zane Jaunmuktane, Juan Eugenio Iglesias

## Abstract

Structural brain analysis at the subregion level offers critical insights into healthy aging and neurodegenerative diseases. The NextBrain histological atlas was recently introduced to support such fine-grained investigations, but its existing Bayesian segmentation framework remains computationally prohibitive, particularly for large-scale studies. We present a new, open-source tool that dramatically accelerates segmentation using a hybrid approach combining: machine learning, contrast-adaptive segmentation; target-specific image synthesis; and fast diffeomorphic registration (all three with GPU support). Our method enables highly granular segmentation of brain MRI scans of any resolution and contrast (*in vivo or ex vivo*) at a fraction of the computational cost of the original method (*<*5 minutes on a GPU). We validate our tool on four different modalities (*in vivo* MRI, *ex vivo* MRI, HiP-CT, and photography) across a total of approximately 4,000 brain scans. Our results demonstrate that the accelerated approach achieves comparable accuracy to the original method in terms of Dice scores, while reducing runtime by over an order of magnitude. This work enables high-resolution anatomical analysis at unprecedented scale and flexibility, providing a practical solution for large neuroimaging studies. Our tool is publicly available in FreeSurfer (https://surfer.nmr.mgh.harvard.edu/fswiki/HistoAtlasSegmentation).

## 1 Introduction

Image segmentation is a fundamental component of human neuroimaging. Automatic delineation of anatomical structures on MRI scans is critical for quantifying volumes of regions of interest (ROIs), assessing their morphometry, and supporting downstream analyses such as functional localization or connectivity modeling. Structural segmentation facilitates the study of brain development (Giedd et al., 1999; Gogtay et al., 2004), aging (Bethlehem et al., 2022), and various neurological and psychiatric disorders (Jack Jr et al., 1999), and is routinely used to derive imaging biomarkers for conditions such as Alzheimer’s disease, schizophrenia, and multiple sclerosis (Cuingnet et al., 2011; Dickerson et al., 2011; Frisoni et al., 2010).

As the scale and diversity of neuroimaging datasets increase, robust and generalizable segmentation tools are needed to handle variation in acquisition protocols, scanner vendors, and patient populations. This is particularly important when sharing open-source tools (as opposed to developing in-house solutions), since few assumptions can be made on the MRI contrast properties of the input scans. For this reason, Bayesian segmentation methods based on generative models (Ashburner & Friston, 2005; Pohl et al., 2006; Puonti et al., 2016; Van Leemput et al., 2002; Wells et al., 1996; Zhang et al., 2002) have long been central to widespread neuroimaging software pipelines such as SPM (Friston et al., 1994), FreeSurfer (Fischl, 2012), FSL (Smith et al., 2004), or AFNI (Cox, 1996).

In terms of generalizability, a particularly interesting subclass of Bayesian segmentation methods uses unsupervised likelihood models, which make it adaptive to MRI contrast. Best represented by Unified Segmentation (Ashburner & Friston, 2005) (distributed with SPM), these methods combine supervised anatomical priors (probabilistic atlases of anatomy) with unsupervised likelihoods that typically comprise a Gaussian mixture model (GMM) and a bias field model. Crucially, the parameters of the GMM and the bias field model are estimated directly from the input scan, which can thus have been acquired with any pulse sequence. Furthermore, because the models of anatomy and image formation are decoupled, high-resolution *ex vivo* data (e.g., histology, *ex vivo* MRI) can be leveraged to construct detailed atlases, which can subsequently be applied to the segmentation of markedly different, lower-quality *in vivo* images. We have successfully used *ex vivo* MRI and histology to build probabilistic atlases of the hippocampus (Iglesias et al., 2015), amygdala (Saygin et al., 2017), and thalamus (Iglesias et al., 2018) at the subregion level, which can be flexibly applied to *in vivo* segmentation.

Our group has recently extended *ex vivo* atlasing from a couple of brain structures to the whole human brain using histology (Casamitjana et al., 2025). Our new atlas (“NextBrain”) comprises 333 ROIs per hemisphere, defined at 200 µm resolution. NextBrain includes a Bayesian segmentation framework capable of transferring this anatomical detail to *in vivo* MRI images. However, the original implementation is computationally intensive: segmenting a single scan at 300 µm resolution can take *days* on a standard CPU-based workstation. While GPU acceleration can reduce run time, it requires access to high-end hardware that is not available to many users, limiting the practicality of the tool for large-scale studies or clinical applications.

One potential solution to this problem would naturally be deep learning, which provides fast inference times and impressive results in a variety of medical image segmentation tasks (Isensee et al., 2021; Ronneberger et al., 2015), including neuroimaging (Henschel et al., 2020; Pereira et al., 2016; Salehi et al., 2017)). Compared with Bayesian segmentation, these approaches suffer from “domain shift”, i.e., they often struggle to generalize to unseen domains without careful finetuning or domain adaptation (Guan & Liu, 2021; Wang & Deng, 2018). There have been attempts to address this domain shift problem; one representative example for brain MRI segmentation is SynthSeg (Billot et al., 2023), which relies on domain randomization. Its design is directly inspired by Bayesian segmentation, and it relies on generating synthetic images with almost the same generative model. Importantly, these images are synthesized with random model parameters to simulate broad variations in image appearance, which leads to high robustness at test time. Despite these advances, current deep learning approaches remain limited in transferring high-resolution information from *ex vivo* domains to *in vivo* data, particularly when only sparse ground-truth annotations are available across a large number of ROIs – as in NextBrain.

In this work, we present a fast and robust reimplementation of Bayesian segmentation that enables segmentation with NextBrain in practical times without specialized hardware. Our method leverages carefully chosen approximations, recent advances in deep learning for initialization, contrast-adaptive synthetic image generation, and fast diffeomorphic registration to deliver highly detailed segmentations of *in vivo* and *ex vivo* brain MR scans with minimal computational demands. Crucially, the pipeline runs without any manual parameter tuning and achieves performance comparable to the original implementation, while reducing run time by more than an order of magnitude. This contribution makes detailed brain parcellation accessible to a wider community and enables high-resolution neuroanatomical analysis at scale.

## 2 Methods

### 2.1 Bayesian segmentation and NextBrain

#### Preliminaries

Bayesian segmentation is typically formulated as an inference problem within a generative model of the data:

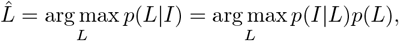

where *L* denotes the label assignment (i.e., the segmentation), and *I* represents the observed image. In general, both the likelihood *p*(*I*|*L*) and the prior *p*(*L*) are governed by parameter sets, denoted *θ*_*I*_ and *θ*_*L*_, respectively. Exact posterior inference requires marginalization over these parameters, which is intractable in practice. A common approximation is to compute point estimates of the parameters by maximizing their posterior distribution:

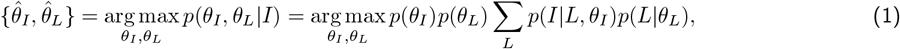

where we have made the standard assumption that the priors over the two parameter sets are independent, i.e., *p*(*θ*_*I*_, *θ*_*L*_) = *p*(*θ*_*I*_)*p*(*θ*_*L*_). Given the estimated parameters, the final segmentation is computed by maximizing the posterior over labelings:

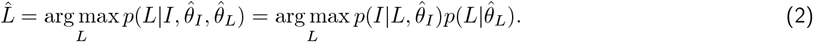

### Model Instantiation in NextBrain

NextBrain (Casamitjana et al., 2025) uses a generative model similar in structure to Unified Segmentation (Ashburner & Friston, 2005) and SAMSEG (Puonti et al., 2016), which we briefly summarize here. The prior is defined by a probabilistic atlas *A*, which specifies conditionally independent categorical distributions over *K* possible labels at each spatial location. This atlas is deformed using a nonlinear transformation ***ϕ***, parameterized by *θ*_*L*_. The deformation follows a probabilistic model encoded by a prior distribution *p*(*θ*_*L*_), which penalizes excessive geometric distortion via a membrane energy term. Specifically, the probability distribution over the labeling *L* is defined as:

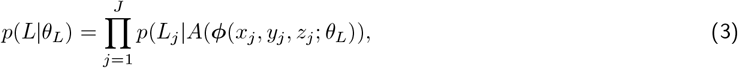

where *j* indexes the *J* voxels in the image, and (*x*_*j*_, *y*_*j*_, *z*_*j*_) represents the spatial coordinates of voxel *j*. The prior over the deformation parameters *θ*_*L*_ is given by:

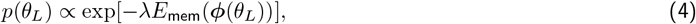

where *λ* is a regularization constant, and *E*_mem_ denotes the membrane energy, defined as:

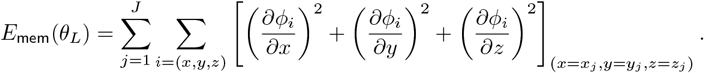

Given a labeling *L*, the likelihood models the image intensities using a GMM, assuming conditional independence across voxels and incorporating a smoothly varying multiplicative bias field. To simplify the model, the image intensities *I* are log-transformed, so that the bias becomes additive and the need to rescale probability densities due to multiplicative bias is eliminated. The likelihood is given by:

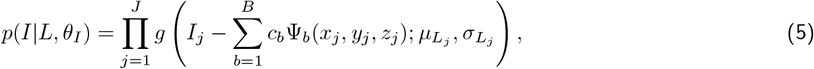

where *g*(*·*; *µ, σ*) denotes the probability density function of a Gaussian distribution with mean *µ* and standard deviation *σ*, and 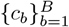 are the coefficients of the bias field basis functions 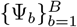. The full set of likelihood parameters is given by *θ*_*I*_ = *{µ*_1_, …, *µ*_*K*_, *σ*_1_, …, *σ*_*K*_, *c*_1_, …, *c*_*B*_*}*, which includes class-specific means and variances, along with the bias field coefficients. A non-informative (uniform) prior is assumed over *θ*_*I*_, such that no specific MRI contrast of bias field is favored: *p*(*θ*_*I*_) *∝* 1.

### Segmentation as Bayesian inference

Substituting the prior and likelihood models (Equations 3, 4, and 5) into Equation 1, taking logarithm, and disregarding constant terms that do not affect the optimization, the following objective function is obtained:

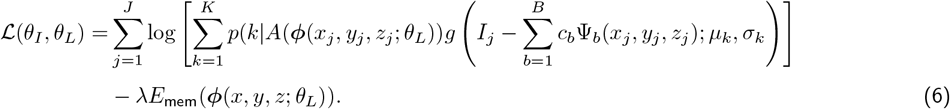

This objective function is maximized using a coordinate ascent scheme, in which the parameters *θ*_*L*_ and *θ*_*I*_ are alternately optimized. The prior parameters *θ*_*L*_ (representing the atlas deformation) are updated using standard numerical optimization methods (Nocedal & Wright, 2006). The likelihood parameters *θ*_*I*_ are optimized using the generalized Expectation Maximization (GEM) algorithm (Dempster et al., 1977). GEM alternates between constructing a lower bound on the objective ℒ via Jensen’s inequality, and maximizing this bound with respect to *θ*_*I*_. This iterative process guarantees an increasing sequence of values for the original objective ℒ.

Maximizing the lower bound involves iteratively updating the means, variances, and bias field parameters. All of these can be updated in closed form given the others. After convergence of this coordinate ascent procedure, the estimated parameters 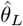 and 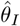 are used to compute the final segmentation, as described in Equation 2. In practice, explicit computation of this equation is unnecessary, since voxel-wise posterior probabilities over labels (also known as “responsibilities”) are already computed as part of the GEM algorithm. These responsibilities are used in the E-step to construct the lower bound. For further details on this framework, the reader is referred to Ashburner and Friston, 2005 or Puonti et al., 2016.

### Computational bottlenecks

The original inference algorithm in NextBrain is computationally intensive and often impractical for routine use. Its complexity arises from a deeply nested optimization structure at three levels: *(i)* alternating between deforming the atlas and updating intensity model parameters; *(ii)* the latter uses a GEM algorithm that iterates between E and M steps; and *(iii)* the M-step itself involves iterative updates of class means, variances, and bias field coefficients, each depending on the others.

NextBrain’s nested optimization structure results in long run times, especially when combined with its large number of anatomical labels and high spatial resolution. NextBrain includes 333 ROIs per hemisphere, which are coarsely grouped during inference but ultimately expanded to the full label set when computing the final segmentation. The atlas is defined at 0.2 mm isotropic resolution, i.e., 125 times more voxels than a standard 1 mm MRI. Even at the default working resolution of 0.3 mm, voxels are 37 times smaller. Although resolution could be further reduced to alleviate computational requirements, this would compromise the benefits of a high-resolution histological atlas.

The combination of model complexity, label granularity, and high spatial resolution results in substantial computational and memory demands. Segmenting a single case at 0.3 mm resolution takes over 48 hours on a CPU and still requires about 60 minutes on a GPU with at least 24 GB of memory. *Ex vivo* segmentation at 0.2–0.3 mm resolution is often infeasible with the current implementation. This large computational footprint limits the practicality of NextBrain as a segmentation tool. In the next section, we present a reimplementation that preserves the advantages of using our high-resolution atlas while dramatically improving run time and hardware efficiency through targeted approximations and algorithmic redesign.

### 2.2 Approximate inference for fast segmentation

#### Overview

Here we present a fast segmentation method for NextBrain that efficiently approximates the inference problem defined in Equation 1. Unlike traditional approaches, our algorithm visits each set of model parameters only once, using GPU-enabled machine learning and a custom GPU-accelerated diffeomorphic registration algorithm. Importantly, the method preserves robustness to variations in MRI modality and contrast by maintaining contrast-adaptive modeling of image intensities. Although the registration involves several hyperparameters, our publicly available implementation uses a fixed set of defaults (employed in all experiments) that do not require manual tuning.

#### Preprocessing

The input scan is first segmented using the segmentation module of our foundation model “BrainFM” (Liu et al., 2025). This method closely follows SynthSeg (Billot et al., 2023), and produces coarse segmentations at 1 mm resolution that are highly robust to variations in contrast, resolution, and imaging artifacts (including strong bias fields). Compared to the original SynthSeg model, BrainFM adds several labels, including five contralateral ROIs in the limbic system (Greve et al., 2021) and 12 extracerebral structures (see Liu et al., 2025 for details).

These coarse segmentations are used to exclude non-brain tissue, including cerebrospinal fluid (CSF), which NextBrain models with the same class as the background. The segmentation is also used to divide the brain into hemispheres. Non-hemispheric ROIs such as the brainstem and white matter lesions are also split by thresholding the left-right component of a deformation field mapping the scan to a symmetric MNI atlas; this deformation is computed within BrainFM using the Registration-by-Regression (RbR) algorithm (Gopinath et al., 2024; Hu et al., 2025). In addition to preprocessing, the ROI segmentations also play a key role when estimating model parameters in later stages of the pipeline, as described below.

#### Optimizing the bias field parameters *{c*_*b*_*}*

In practice, Bayesian segmentation algorithms group ROIs into a smaller number of tissue classes during inference, clustering regions expected to share similar intensity profiles (e.g., gray matter structures such as the cerebral cortex, hippocampus, and amygdala). This strategy improves robustness, especially for small regions that may otherwise yield noisy parameter estimates. For instance, since the third ventricle is a thin layer of cerebrospinal fluid, its intensity distribution can be more reliably estimated when grouped with the much larger lateral ventricles. Grouping also benefits bias field estimation by signaling that some ROIs should have similar intensity profiles, despite potentially being distant from each other (e.g., left and right putamen); this information greatly helps disambiguate intensity variations due to bias field and anatomy.

In our proposed method, we model the bias field with a 3D discrete cosine transform (DCT) basis of order up to 6, resulting in 343 basis functions. To estimate the bias field parameters *{c*_*b*_*}*, we first extract the whole brain using the available segmentation and group the BrainFM labels into eight tissue classes: cerebral gray matter, cerebellar gray matter, cerebral white matter, cerebellar white matter, brainstem, cerebrospinal fluid (CSF), thalamus, and pallidum. A soft segmentation is then generated by blurring the one-hot encoding of this 8-class map with a Gaussian kernel (*σ* = 0.5 *mm*). This soft segmentation is used to perform a *single M-step* of the GEM algorithm, with responsibilities fixed to the soft assignments. Due to the high reliability of the SynthSeg-based segmentation, the large number of voxels available, and the relatively low number of parameters, reclassifying voxels in an EM loop offers negligible benefit. While the M-step entails a nested optimization that alternates between estimating tissue class statistics (means and variances) and updating the bias field coefficients, convergence is rapid.

#### Optimizing the prior parameters (atlas registration) *θ*_*L*_

In Bayesian segmentation, updating the atlas deformation corresponds to optimizing the objective in Equation 6 with respect to the deformation parameters *θ*_*L*_, while holding the intensity model parameters *θ*_*I*_ fixed. This step amounts to a deformable image registration problem, where the data term is given by the log-likelihood of the image under the atlas and the current estimate of *θ*_*I*_. However, this registration becomes computationally expensive in the classical coordinate ascent scheme, as every *θ*_*I*_ update effectively changes the objective function with respect to *θ*_*L*_, making the optimization slow.

To avoid this costly alternation, we leverage the available BrainFM segmentation to bypass the need to update *θ*_*I*_ iteratively. Despite its lower resolution compared to the NextBrain atlas, the BrainFM segmentation is sufficiently accurate to estimate class-wise means and variances from the bias field corrected image. We use these parameters to synthesize a Gaussian image (a “cartoon”) from the probabilistic atlas, which is then resampled to match the resolution of the input scan (Figure 1). This procedure ensures that the cartoon exhibits contrast and resolution closely resembling those of the target scan, thereby greatly facilitating registration in the subsequent steps described below.

**Figure 1:**
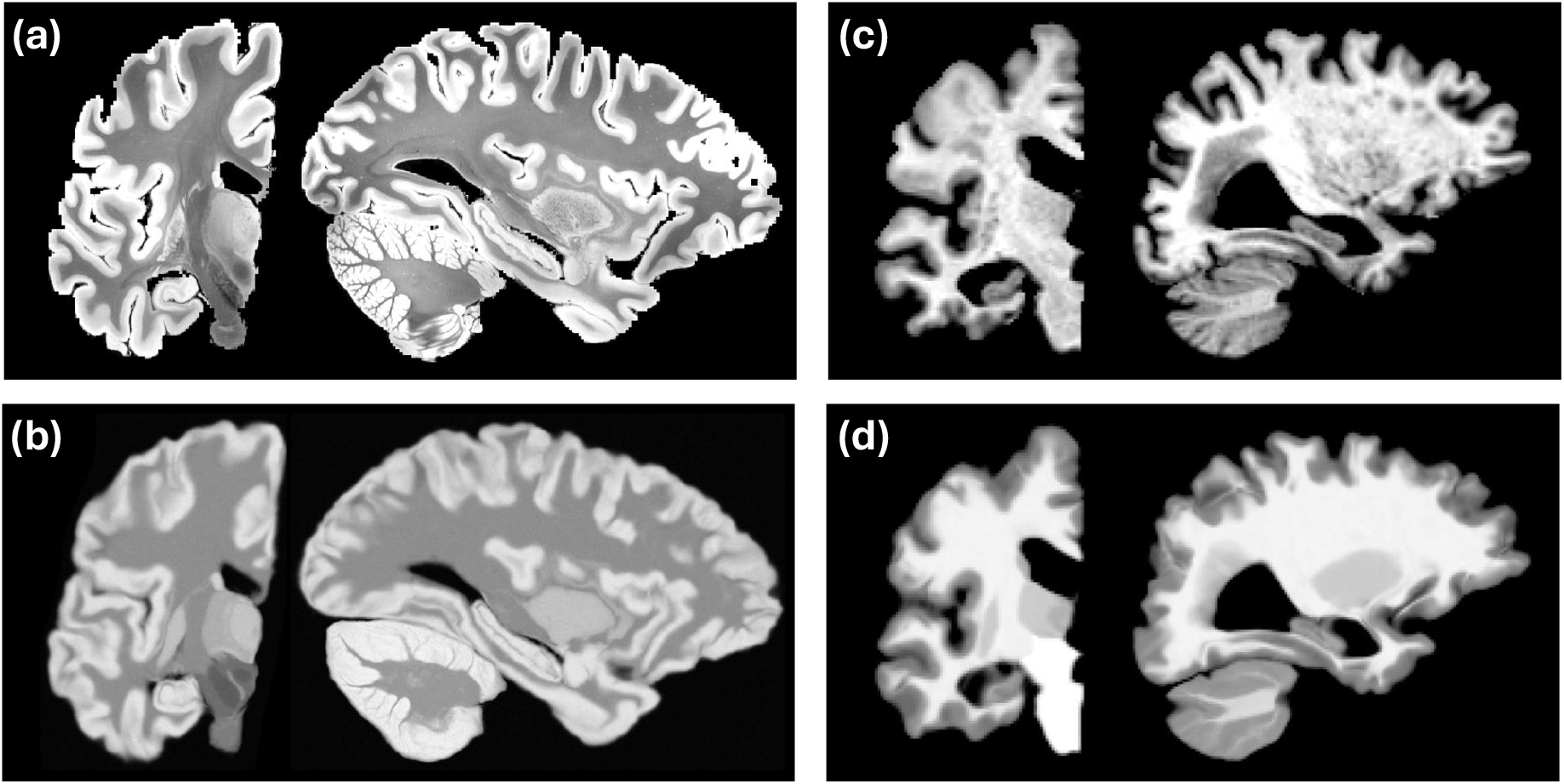
(a) Coronal and sagittal slices of an *ex vivo* scan at 0.2 mm resolution (Edlow et al., 2019). (b) Matching cartoon after registration with FireANTs and composite similarity metric. (c) Slices of a 1 mm isotropic, skull-stripped, T1-weighted scan from the MIRIAD dataset (Malone et al., 2013). (d) Registered cartoon. Note the differences in contrast and resolutions of the cartoons (c,d), which attempt to match the scans they seek to segment.

Because the NextBrain atlas includes a large number of ROIs, we group them into 16 tissue classes to generate a sufficiently detailed cartoon: cerebral white matter, cerebral gray matter, cerebellar white matter, cerebellar cortex, caudate, putamen, pallidum, lateral thalamus, medial thalamus, red nucleus, compact brainstem white matter, diffuse brainstem white matter, hypothalamus, mammillary bodies, dentate nucleus of the cerebellum, and hippocampal white matter. The mean intensities for these groups are derived from the BrainFM segmentation when a direct label correspondence exists (e.g., cerebral white matter); otherwise, we use heuristic estimates based on linear combinations of gray and white matter means (see Supplementary Material for details).

We perform registration between the synthetic cartoon and the input image one hemisphere at a time, as NextBrain is a single-hemisphere model. Registration is performed using FireANTs (Jena et al., 2024), which we run in greedy, non-symmetric mode (i.e., with the loss computed in the fixed image grid). The similarity metric comprises two terms: *(i)* local normalized cross-correlation (LNCC, Avants et al. 2008), computed with a kernel radius of 3 voxels; and *(ii)* the mean Dice score between the BrainFM segmentation and the warped atlas labels, where the atlas ROIs are grouped to match the BrainFM protocol. The optimization is carried out using the Adam optimizer (Kingma & Ba, 2014), and proceeds in a multiscale fashion, from coarse to fine resolution levels.

While FireANTs does not explicitly include a regularization term in the objective function, smoothness is implicitly enforced through the multiscale optimization scheme and the use of diffeomorphic transformations parameterized in a smooth velocity field space. This setup discourages implausible deformations by naturally limiting high-frequency updates, ensuring anatomically consistent mappings without the need for hand-tuned regularization weights. We used Gaussian kernels with *σ* = 0.25 and *σ* = 1.0 to smooth the field and its gradient, respectively. Examples of brain scans of different contrast and resolution and their corresponding registered cartoons are shown in Figure 1.

#### Optimizing the Gaussian parameters 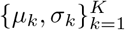 and obtaining the final segmentation

Given the estimated bias field and atlas deformation parameters, the class means and variances can be optimized using the EM algorithm; we note that it is EM and not GEM because the fixed bias field enables global maximization of the lower bound at each iteration. The EM procedure is carried out at a predefined resolution, to which both the input image and the probabilistic atlas are resampled. In our implementation, we set this resolution to 0.4 mm for *in vivo* scans, which balances anatomical detail and computational efficiency. For *ex vivo* scans, we set it to 0.2 mm (the resolution of the atlas) or the resolution of the scan, whichever is lower.

EM inference is performed over the same 16 tissue classes used previously to generate the cartoon for registration. Consequently, the soft assignments (responsibilities) from the final E-step yield a probabilistic segmentation into 16 tissue classes rather than fine-grained ROIs. Posterior probabilities over the full set of ROIs are obtained by assigning each ROI a fraction of the posterior equal to the ratio between its prior probability and that of the associated tissue class (Puonti et al., 2016).

### Other simplifications and practical details

To further improve the efficiency and practicality of our method, we implemented some additional algorithmic and engineering simplifications:

- As mentioned above, we segment images at 0.4 mm isotropic resolution, instead of the 0.3 mm resolution used in the original NextBrain implementation. This reduces computational and memory demands by a factor of 2.4 while preserving anatomical detail.
- The entire pipeline, including FireANTs, is implemented in PyTorch, enabling seamless deployment on both GPU and CPU. PyTorch also facilitates efficient multi-threaded CPU execution.
- To accelerate atlas deformation, we store the probabilistic atlas as a set of sparse vectors defined within tight bounding boxes for each ROI, which allows rapid deformation of only the relevant subregions.
- In practice, GPU memory usage is dominated by the estimation of likelihood parameters, whereas actual computation is dominated by the atlas registration. When processing high-resolution *ex vivo* images at resolutions finer than 0.4 mm, we provide the option to offload likelihood optimization to the CPU (where memory is typically more abundant) while keeping the registration step on the GPU.
- To further reduce run time and memory usage, we simplified the original NextBrain atlas by removing the smallest ROIs. The final model includes 264 anatomical regions instead of 333; the list is provided in Table S2.

On modern hardware, a full case (both hemispheres) processes in approximately 5 minutes on a GPU, or 20 minutes on a multi-core CPU workstation.

## 3 Experiments and Results

### 3.1 Data

We use five different datasets to evaluate the proposed method:

### Ex vivo MRI

This dataset consists of 21 specimens imaged with a Siemens 7 Tesla scanner at 0.12 mm isotropic resolution. The average age of the cohort is 65 years and the samples are from donors without neurological disorders. The samples comprise 12 single left hemispheres, 7 single right hemispheres, and two full brains. One of the full brains corresponds to the case described in Edlow et al., 2019, for which we manually annotated 98 ROIs (including subcortical regions) in the right hemisphere at 0.2 mm isotropic resolution; these labels were released as part of the NextBrain dataset (Casamitjana et al., 2025). No gold-standard segmentations are available for the rest of the cases.

### In vivo MRI (OpenBHB)

OpenBHB (Dufumier et al., 2022) is a public meta-dataset with *∼*1 mm isotropic T1-weighted volumes of 3,227 healthy subjects scanned at over 60 sites. The age range of the subjects is 6-86 years (mean: 25.2 years). No ground-truth segmentations are available for these scans.

### In vivo MRI (MIRIAD)

The Minimal Interval Resonance Imaging in Alzheimer’s Disease (MIRIAD) (Malone et al., 2013) is a public dataset with *∼*1 mm isotropic T1-weighted volumes of 69 subjects (46 mild–moderate Alzheimer’s subjects and 23 controls, age at entry 69.5 *±* 7.1 years, 31 men). It comprises 708 scans acquired with the same scanner and pulse sequences at intervals of 2, 6, 14, 26, 38 and 52 weeks, 18 and 24 months from baseline, with up to 12 scans per individual. Out of the 708 scans, 185 are repeat acquisitions of the same timepoints, which can be used to assess test-retest reliability. No ground-truth segmentations are available for these scans.

### Ex vivo Hierarchical Phase-Contrast Tomography (HiP-CT)

HiP-CT is an advanced X-ray imaging technique used to visualize intact human organs *ex vivo* at multiple scales, from whole-organ structure down to cellular detail (Walsh et al., 2021). It leverages phase contrast generated by X-ray wavefront distortions using a synchrotron source to achieve resolutions as high as 1 *µ*m. Walsh et al., 2021 have made 6 brain HiP-CT scans available (4 males, 2 females, ages 63-80). The scans were downloaded from the Human Organ Atlas (HOA) repository at a resolution that was closest to 0.3 mm isotropic, which resulted in a range of resolutions spanning from 0.2–0.3 mm. We excluded case LADAF-2020-31, as it was among the earlier acquisitions and its image quality was notably inferior to that of the other cases. No ground-truth segmentations are available for these scans.

### Ex vivo Visible Human 2.0

This is a high-resolution anatomical dataset of the head and neck of a 66-year-old male donor, focused on axial cryosection photography acquired every 0.15 mm (Ratiu et al., 2003). Tissue contrast was optimized to distinguish fine brain structures. We normalized the photographs by median intensity to correct for illumination inhomogeneity across slices.

### 3.2 Results

#### 3.2.1 Segmentation of ultra-high resolution ex vivo MRI

Figure 2 presents segmented slices from the the case described in Edlow et al., 2019, including both the manually labeled ROIs and the corresponding automated segmentations. Despite the difference in granularity between the two approaches (98 manually defined ROIs versus 264 in the automated output), the overall agreement between them is remarkably strong. Table S1 in the Supplementary Material summarizes the Dice scores for the 98 manually labeled ROIs, obtained by clustering the automated segmentation to align with the gold standard definitions. ROI sizes are also included in the table. As expected, there is a clear association between region size and Dice score. Larger structures (e.g., cerebral and cerebellar white matter and cortex) achieve Dice values near 0.9. Smaller regions tend to show lower scores, although only a handful fall below 0.4, which remains adequate for localization purposes. Overall, the Dice performance (0.610 *±* 0.207) closely matches that of the original method (0.626 *±* 0.180), indicating comparable segmentation accuracy.

**Figure 2:**
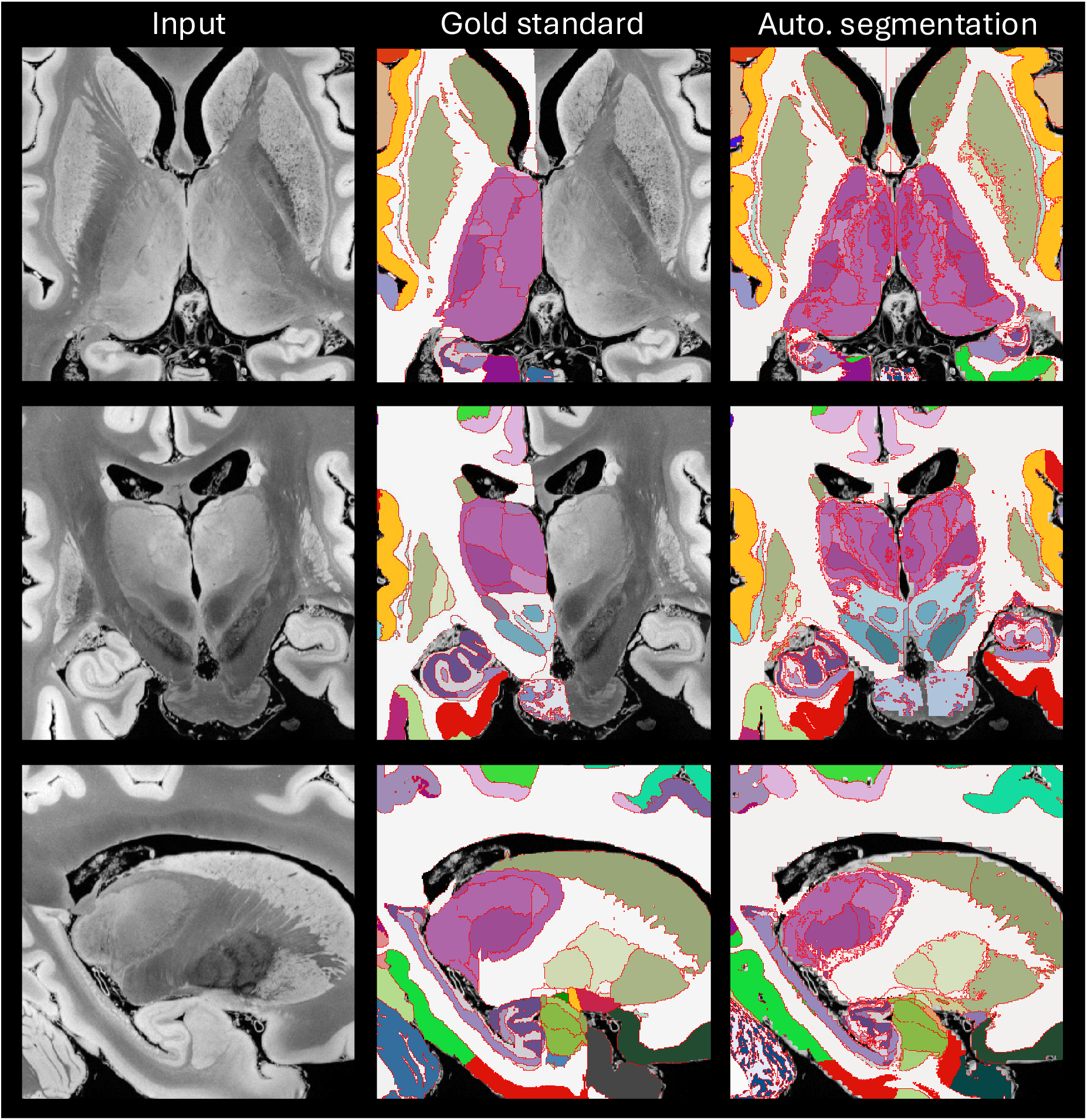
Axial (top row), coronal (middle), and sagittal slices (bottom) of the publicly available *ex vivo* scan from Edlow et al., 2019, along with manual delineations (only right hemisphere available) and the automated segmentation with NextBrain. We note that the input was resampled from 0.1 mm to 0.2 mm resolution, which is the voxel size of the manual segmentation, and also the native resolution of NextBrain. We also note that the manual delineation is less granular than the automated segmentation (98 ROIs vs 264).

Qualitative results are shown in Figure 3, which displays sample coronal segmentations of five other subjects from the *ex vivo* MRI data set. As no reference segmentations are available for 20 out of the 21 subjects, we instead report the volumes of the substructures of the hippocampus, thalamus and amygdala in tables in Tables 1, 2, and 3, and compare them with those from other existing *ex vivo* atlases (Iglesias et al., 2015, 2018; Saygin et al., 2017). Because these atlases were constructed from manual delineations based on anatomical protocols that differ from those of NextBrain, we mapped each label in the atlases to the closest corresponding NextBrain ROI or set of ROIs. The volumes and ROI correspondences are listed in Tables 1, 2, and 3. In general, the volumes agree quite well, with two exceptions: small or narrow structures like the granular layer of the dentate gyrus in the hippocampus; and the posterior regions of the thalamus, due to discrepacies in manual delineation protocols – in Iglesias et al., 2018, the boundary between mediodorsal and pulvinar regions is more anterior than in NextBrain, and the whole thalamus is also generally larger (6,540 vs 5,970 mm^3^).

**Table 1:** Average volumes of the hippocampal subfields from Iglesias et al., 2015 (N=15) and from the automated NextBrain segmentations on the *ex vivo* MRI data (N=20).

| **Hippocampus** |  |  |  |  |
| --- | --- | --- | --- | --- |
| **Structure** | (Iglesias et al., 2015)) |  | Automated NextBrain segmentations |  |
| | Combined structures | Volume $\mu$ (mm<sup>3</sup>) | Combined structures | Volume $\mu \pm \sigma$ (mm<sup>3</sup>) |
| **Subiculum** | Pre/Post-subiculum<br>Subiculum | 624 | Caudal/uncal/rostral-subiculum | 846.72<br>$\pm 109.67$ |
| **Alveus** | Alveus | 320 | Alveus | 427.61<br>$\pm 94.01$ |
| **CA1** | CA1 | 520 | Stratum-radiatum-caudal/uncal/rostral/-CA1<br>Stratum-pyramidale-caudal/uncal/rostral-CA1<br>Stratum-oriens-caudal/uncal/rostral-CA1 | 873.71<br>$\pm 124.55$ |
| **CA2/3** | CA2/3 | 179 | Stratum-radiatum-caudal/uncal/rostral-CA2<br>Stratum-pyramidale-caudal/uncal/rostral-CA2/CA3<br>Stratum-oriens-caudal/uncal/rostral-CA2/CA3 | 239.48<br>$\pm 45.42$ |
| **CA 4** | CA 4 | 211 | pyramidal-cells-caudal/uncal/rostral-CA4<br>molecular-layer-caudal/uncal/rostral-dentate-gyrus | 348.99<br>$\pm 73.72$ |
| **Granular layer dentate gyrus** | GC-DG | 244 | granular-layer-caudal/uncal/rostral-dentate-gyrus | 54.27<br>$\pm 10.94$ |
| **Hippocampal Amygdala transition area** | HATA | 50 | amygdalohippocampal-transition-area | 14.90<br>$\pm 2.96$ |
| **Fimbria** | Fimbria | 92 | Fimbria | 51.91<br>$\pm 22.48$ |
| **Molecular layer** | Molecular layer | 466 | molecular-layer-caudal/uncal/rostral-dentate-gyrus<br>stratum-moleculare-caudal/uncal/rostral-CA1/CA2/CA3 | 714.08<br>$\pm 135.89$ |

**Table 2:** Average volumes of nuclei of the amygdala from Saygin et al., 2017 (N=10) and from the automated NextBrain segmentations on the *ex vivo* MRI data (N=20).

| **Amygdala** |  |  |  |  |
| --- | --- | --- | --- | --- |
| **Structure** | (Saygin et al., 2017)) |  | Automated NextBrain segmentations |  |
| | Combined structures | Volume $\mu \pm se$ (mm<sup>3</sup>) | Combined structures | Volume $\mu \pm \sigma$ (mm<sup>3</sup>) |
| **Lateral nucleus** | Lateral nucleus | 453<br>$\pm 31.4$ | lateral-nucleus | 393.69<br>$\pm 40.78$ |
| **Basal nucleus** | Basal nucleus | 300.9<br>$\pm 19.2$ | basolateral-nucleus<br>(basal-nucleus) | 447.48<br>$\pm 53.29$ |
| **Central nucleus** | Central nucleus | 32.5<br>$\pm 7$ | medial/lateral-subdivision-central-nucleus | 37.51<br>$\pm 10.22$ |
| **Medial nucleus** | Medial nucleus | 21.8<br>$\pm 5.6$ | rostral-subdivision-medial-nucleus | 39.87<br>$\pm 12.41$ |
| **Cortico-amygdaloid Transition Area** | CAT | 174.8<br>$\pm 17.3$ | amygdalocortical-(corticoamygdaloid)<br>transition-area | 152.18<br>$\pm 25.67$ |
| **Paralaminar nucleus** | Paralaminar nucleus | 31.9<br>$\pm 6.4$ | paralaminar-nucleus | 13.16<br>$\pm 3.49$ |
| **Anterior Amygdala Area** | Anterior Amygdala Area | 39.8<br>$\pm 7.9$ | anterior-amygdaloid-area | 113<br>$\pm 20.75$ |

**Table 3:**
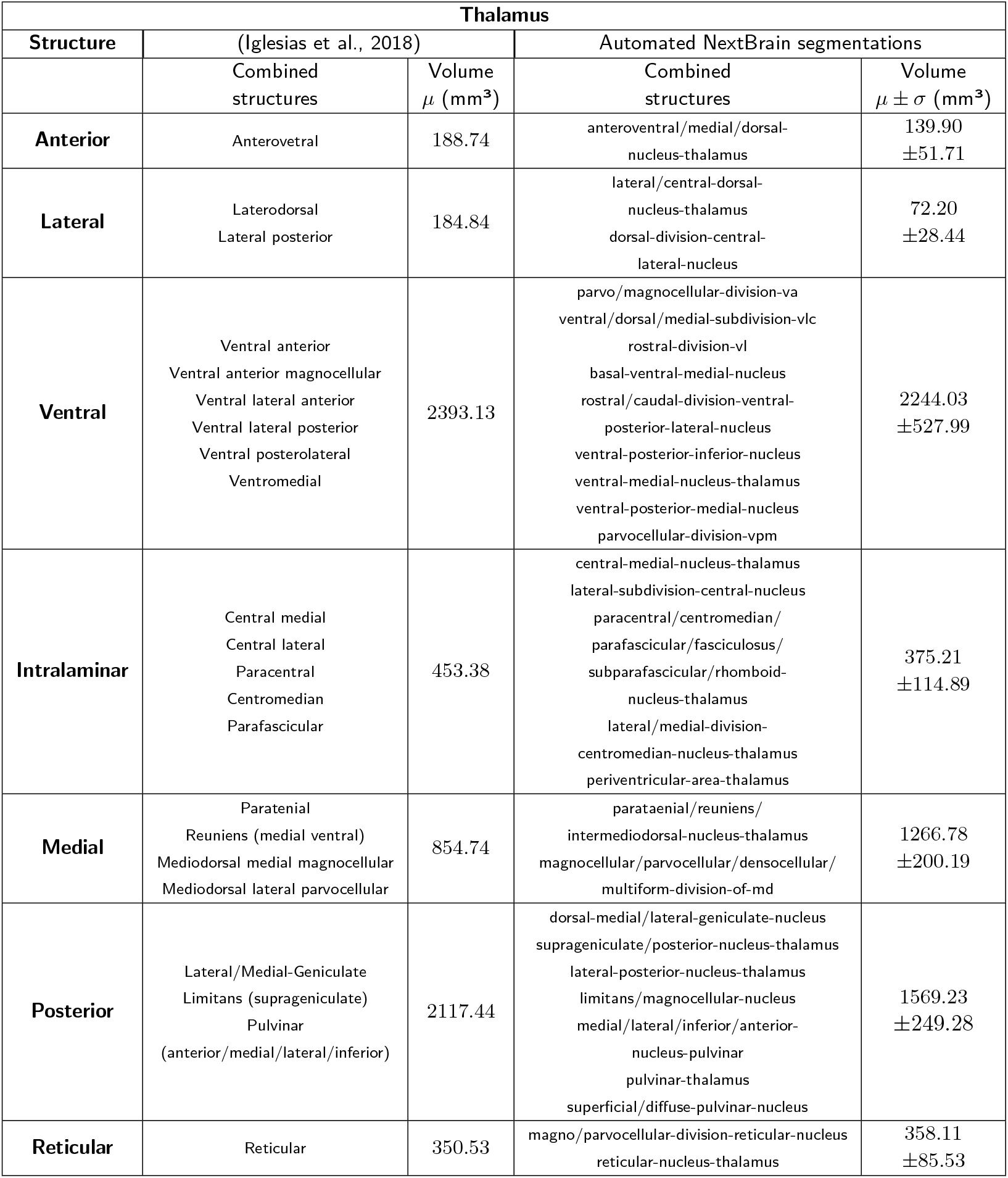
Average volumes of nuclei of the thalamus from Iglesias et al., 2018 (N=6) and from the automated NextBrain segmentations on the *ex vivo* MRI data (N=20).

**Figure 3:**
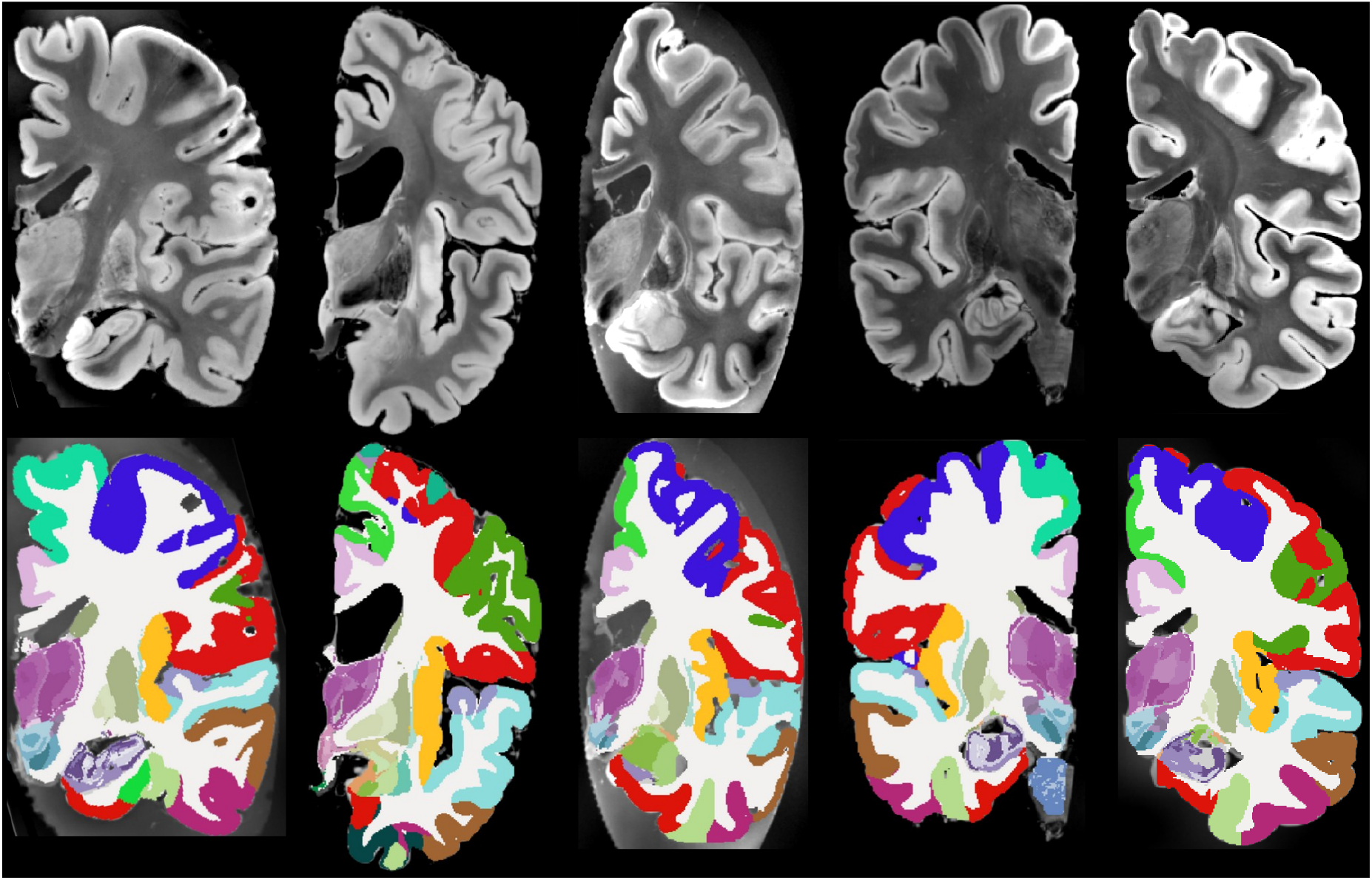
Five example subjects from the ex-vivo MRI data set. Coronal view of the MRI data (top) and the corresponding automated segmentation using NextBrain (bottom).

### 3.2.2 Segmentation of in vivo MRI

#### Dice and volume correlation on OpenBHB

As with most human brain MRI segmentation tools, one of the primary applications of NextBrain is the analysis of *in vivo* scans acquired at near 1 mm isotropic voxel size. However, evaluating subregion-level Dice scores on such data would require manual annotations of structures that often cannot be reliably delineated at this resolution. Instead, we assess performance at the whole-ROI level by merging the automatically segmented subregions to approximate the FreeSurfer protocol (Fischl et al., 2002). As a reference, we used the “supervised baseline” U-Net from Billot et al. (2023), trained on 1 mm isotropic T1-weighted scans, to segment the 3,227 subjects from the OpenBHB dataset. While using automated segmentations as ground truth constitutes a silver (rather than gold) standard, this approach enables evaluation over a much larger and more diverse sample. The heterogeneity of OpenBHB further strengthens the generalizability of the assessment.

Figure 4 illustrates the segmentation results for a representative case from the OpenBHB dataset. Our method produces segmentations that are qualitatively very similar to those of the baseline, while also leveraging visible subregional boundaries and additional anatomical labels to more precisely align the fine-grained NextBrain labels with the underlying image data. Noticeable improvements include: clearer separation between the putamen and adjacent cortex; subdivision of the thalamus that reflects the darker contrast of medial and posterior nuclei; more accurate delineation of the lateral boundary of the thalamus; subdivision of the pallidum; and improved characterization of the hypothalamic and subthalamic regions, among others.

**Figure 4:**
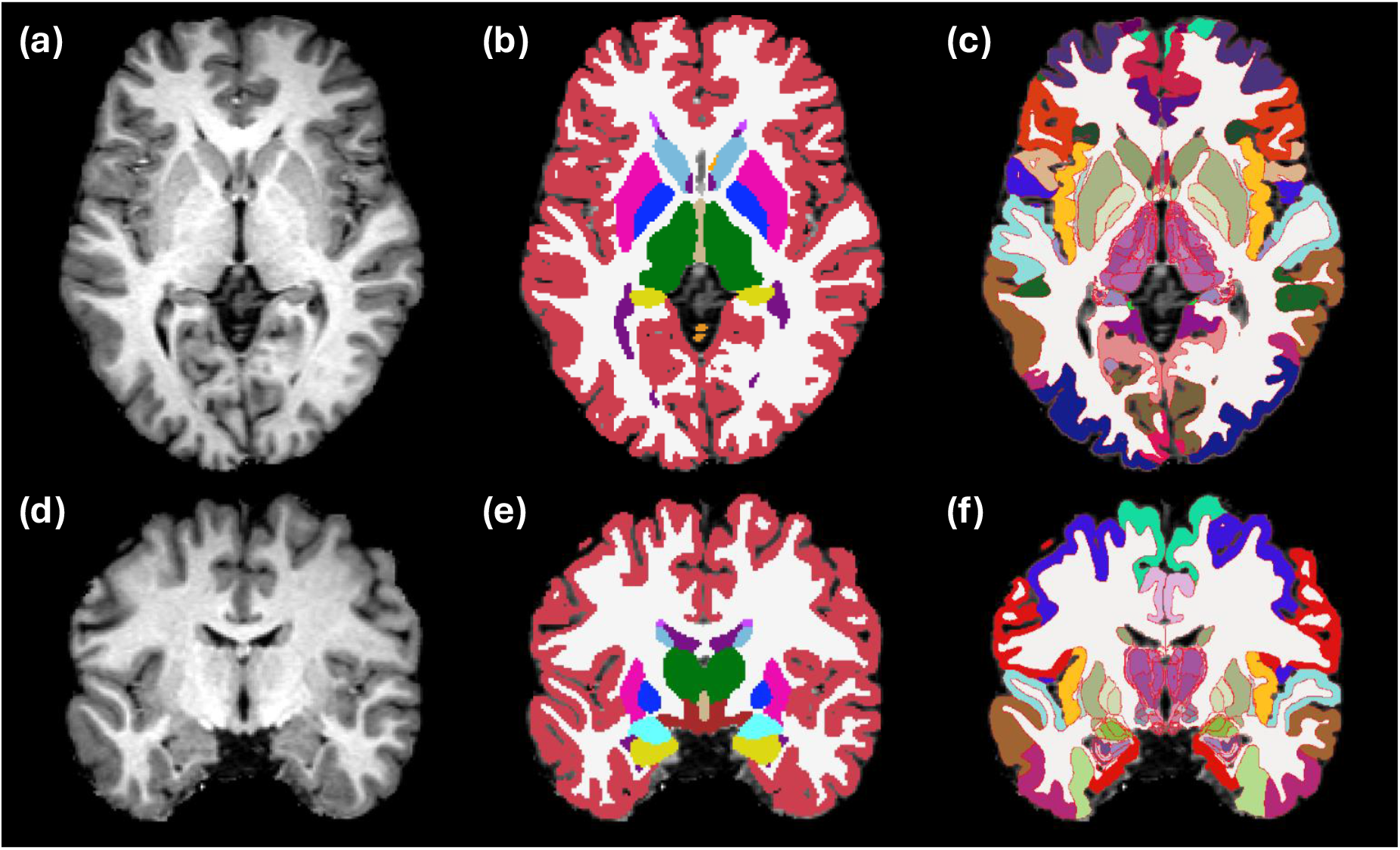
Segmentation of sample subject (368981431172) from OpenBHB dataset. (a) Axial slice of input. (b) Reference segmentation with supervised U-Net for 1 mm T1 scans. (c) Segmentation using NextBrain, with ROI boundaries highlighted in red. (e-f) Coronal slice of the same case.

Table 4 presents the quantitative results for both Dice scores and volume correlations. There is a strong correlation between ROI volumes computed by the two methods. Dice scores comparing the segmentations directly are also consistently high across all regions. These findings support the robustness of the segmentation approach at the whole-region level.

**Table 4:** The average Dice scores and volume correlations against the silver standard on the OpenBHB data set. The NextBrain ROIs are grouped to match the coarser segmentation from the supervised U-Net.

| *ROI* | *Dice* | *Correlation* |
| --- | --- | --- |
| Cerebral white matter | 0.908 | 0.934 |
| Cerebral cortex | 0.865 | 0.920 |
| Cerebellar white matter | 0.837 | 0.861 |
| Cerebellar cortex | 0.910 | 0.909 |
| Thalamus | 0.914 | 0.944 |
| Caudate | 0.894 | 0.816 |
| Putamen | 0.927 | 0.960 |
| Pallidum | 0.894 | 0.899 |
| Hippocampus | 0.881 | 0.867 |
| Amygdala | 0.832 | 0.813 |

Figure 5 illustrates the ROI-wise correlation between age and volume (an “aging map”) based on our high-resolution parcellation. The results closely align with those reported using the original tools in Casamitjana et al., 2025, revealing region-specific patterns of age-related volume change. Notable effects include stronger associations in the frontal cortex, the medial thalamic subregions, the anterior caudate, and the subicular region of the hippocampus.

**Figure 5:**
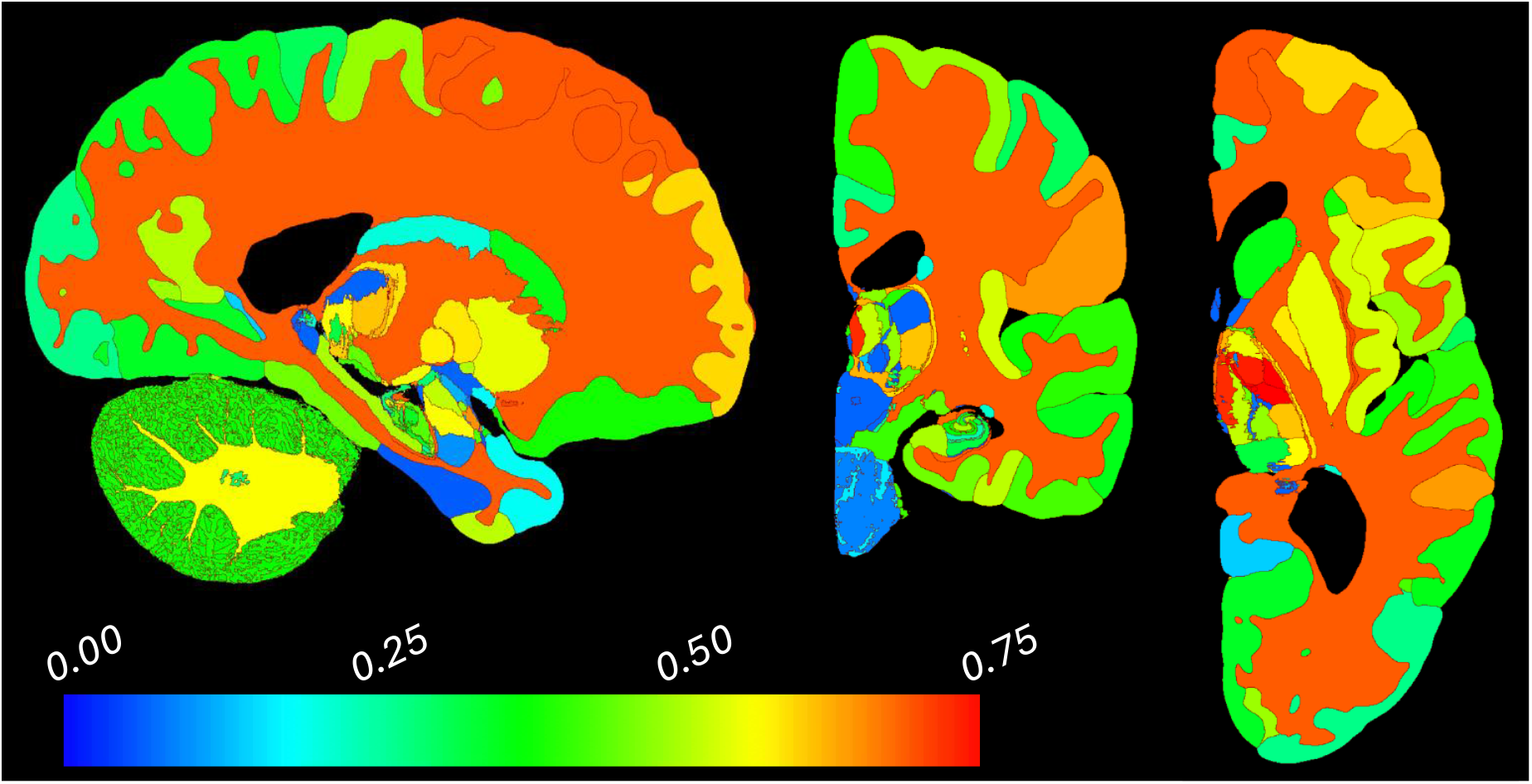
Slices of map of Spearman correlation for ROI volume vs age using OpenBHB dataset, restricted to subjects aged 35 years and over (*n* = 431 subjects, mean age: 57.9 years); we chose this age range as 35 is approximately the age when age-related atrophy begins.

#### Test-retest reliability with MIRIAD

When processing *in vivo* MRI scans at approximately 1 mm resolution, NextBrain necessarily relies more heavily on geometric priors encoded in the atlas to delineate many internal boundaries – some of which are more reliably inferred than others from the outer contours of brain regions. It is therefore important to evaluate the test-retest reliability of the method. For this purpose, we used the MIRIAD dataset, which has 185 timepoints with repeat acquisitions. We computed the intra-class correlation coefficient (ICC) for the volumes of all 264 ROIs, averaged across the left and right hemispheres. An complete list is provided in Table S2 of the supplementary material. The vast majority of ROIs exhibit excellent reliability, with ICC values exceeding 0.8. Notably, all ROIs with lower ICCs are very small in volume (Figure 6): all with ICC*<*0.9 have a volume below 170 mm^3^ (except the myelencephalon), and all with ICC*<*0.8 fall below 30 mm^3^.

**Figure 6:**
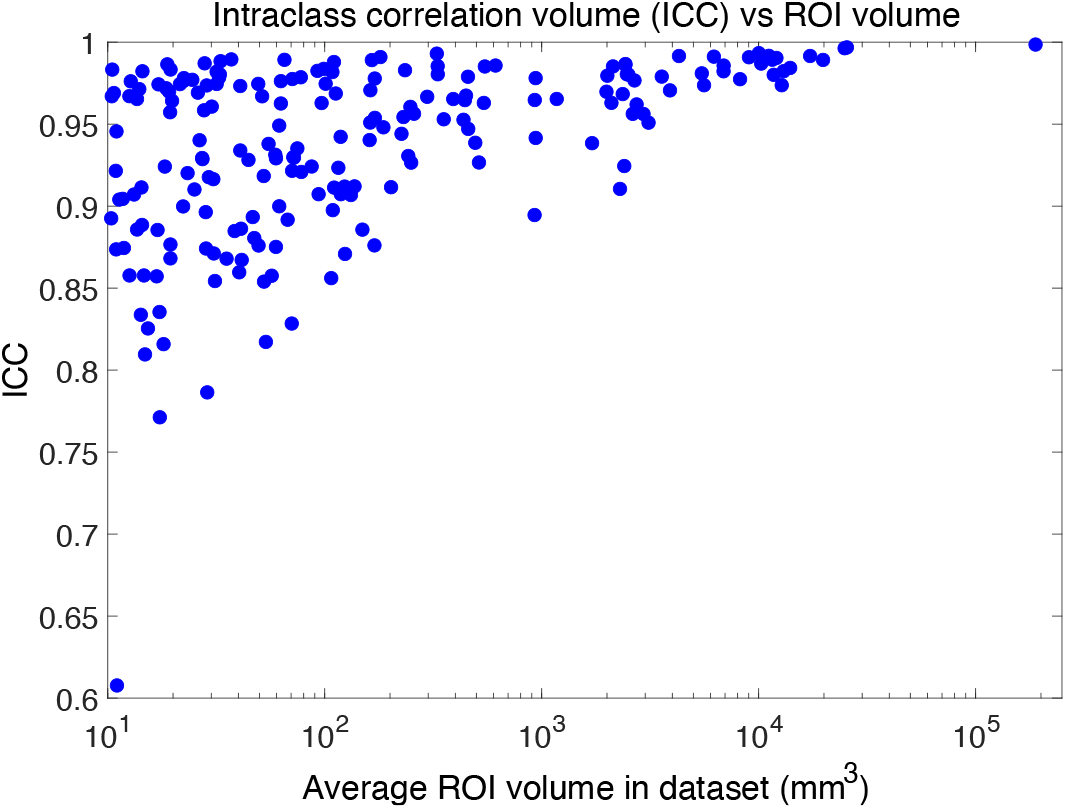
Scatter plot of ICC vs average ROI volume in test-retest experiment on MIRIAD dataset.

#### 3.2.3 Segmentation of unconventional, high-resolution ex vivo modalities

Figure 7 shows coronal slices of all five HiP-CT scans along with the segmentation produced by NextBrain. Notably, our method can tackle significant variations from normal appearing anatomy as exemplified by the three subjects with substantially enlarged or deformed ventricles. Figure 8 shows orthogonal slices from the Visible Human dataset alongside the corresponding segmentation. Given the higher native resolution of this dataset compared to NextBrain, the segmentation was performed at half resolution (0.3 mm isotropic) to reduce computational load. The segmentations were computed on grayscale images; although our method naturally extends to RGB inputs (similar to Bayesian segmentation, with mean vector and convariance matrices replacing scalar means and variances), processing full color volumes at this resolution would require prohibitive amounts of memory. The high-resolution and excellent contrast of both *ex vivo* data sets allow for accurate delineation of complex structures such as the thalamic nuclei. The Visible Human data has increased detail in regions that are often difficult to visualize in MRI, e.g., the dentate gyrus of the cerebellum, as seen in the sagittal slice.

**Figure 7:**
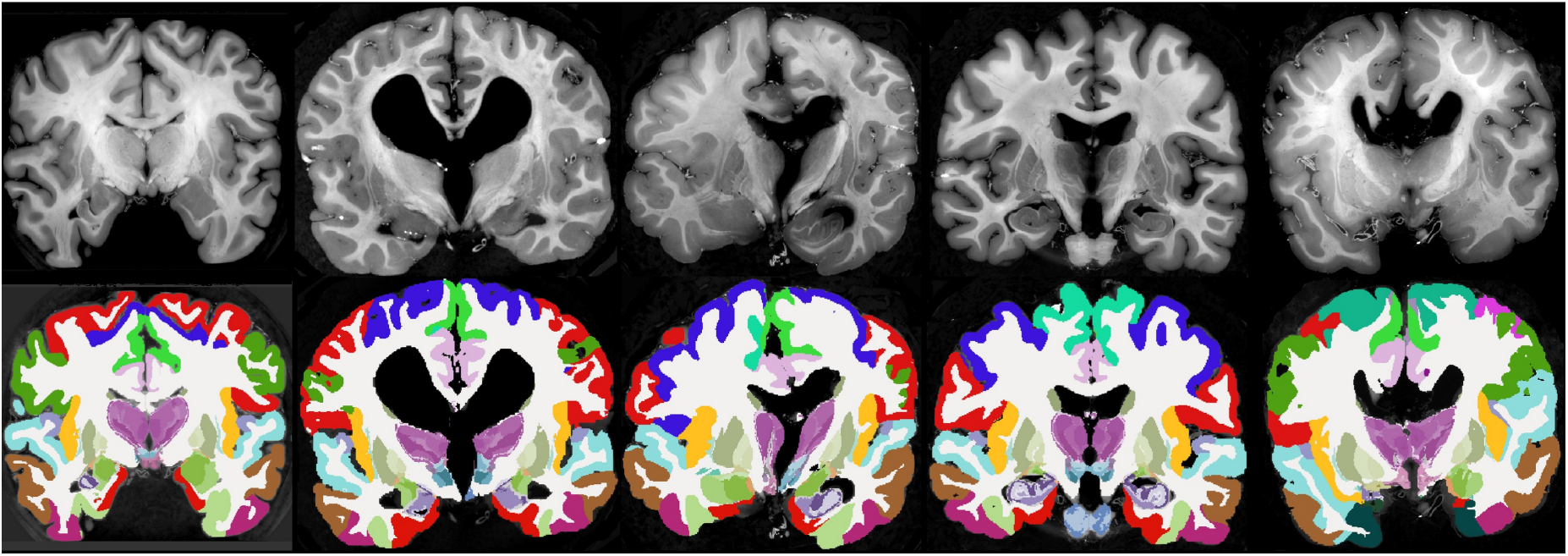
Coronal slices of the five HiP-CT scans (top) and their corresponding automated segmentations using NextBrain (bottom).

**Figure 8:**
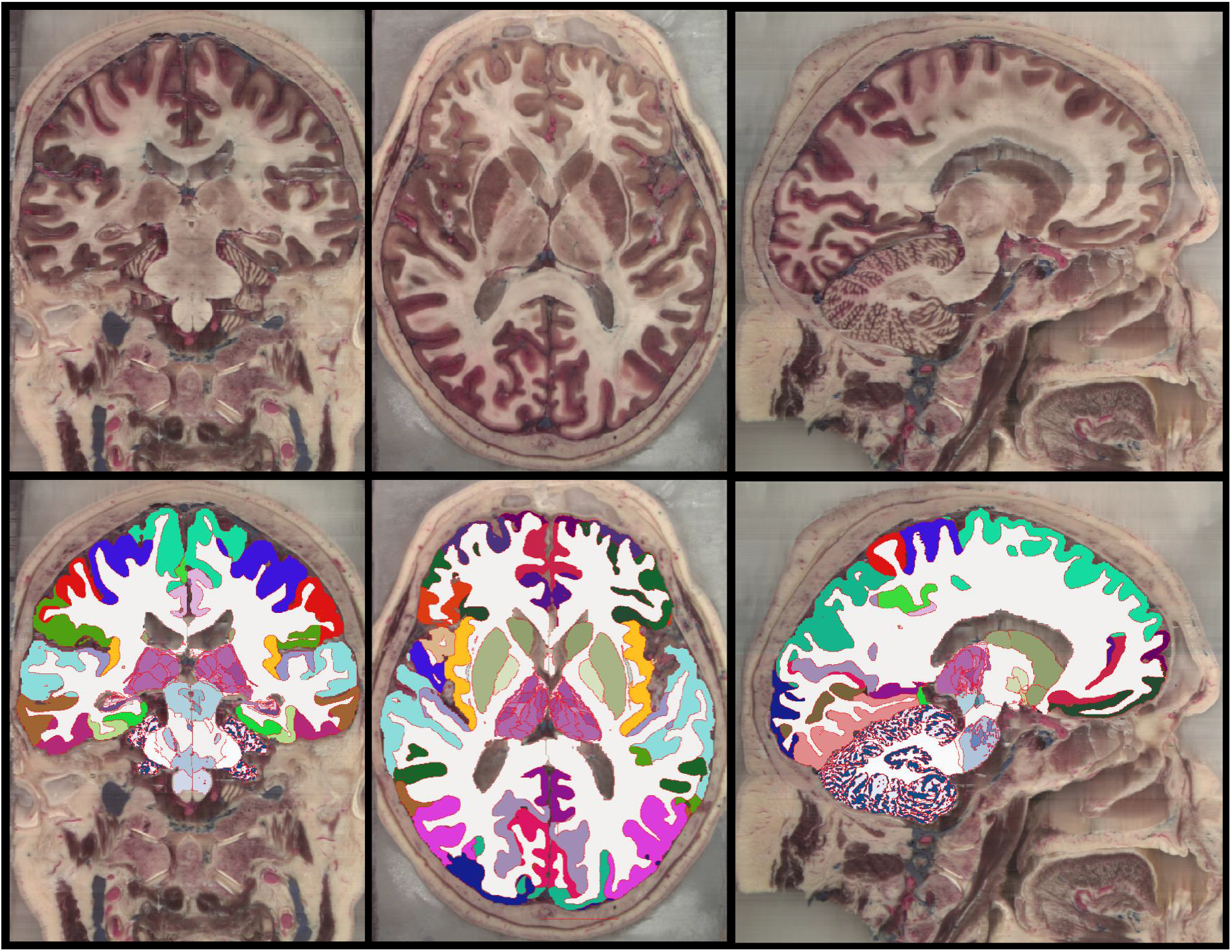
Coronal (left), axial (middle), and sagittal slices (left) of the Visible Human 2.0 dataset, along with automated segmentation computed with NextBrain.

## 4 Discussion and conclusion

In this work, we introduced a fast segmentation method based on the NextBrain histological atlas, capable of accurately parcellating a large number of brain ROIs across several modalities. A key strength of the proposed approach is its computational efficiency, which enables practical application to high-resolution datasets without incurring prohibitive computational costs. For example, segmenting the Visible Human volume at 0.3 mm isotropic resolution required only *∼*90 minutes on a standard eight-core workstation, compared to nearly a week using the original implementation. For *in vivo* scans, our method completes segmentation in approximately 20 minutes on the same hardware, whereas the original code required 2–3 days. When a GPU with sufficient memory (*∼*24 GB) is available, segmentation time can be reduced to under five minutes.

Quantitative validation across several benchmarks, including manual annotations, volume comparisons, and comparison against established segmentation methods, demonstrated that our approach achieves competitive accuracy. As expected, segmentation performance is positively correlated with ROI size, but even small regions such as thalamic subregions, hippocampal subfields, and subregions of the amygdala were reliably identified. Moreover, the method captures biologically meaningful variation: age-volume correlations derived from the segmentations revealed region-specific patterns consistent with previous literature, supporting both the anatomical fidelity and practical relevance of the results.

Two main limitations should be acknowledged. First, our method inherits the constraints of the NextBrain atlas, which was derived from only five cases of older, healthy individuals; consequently, generalization to broader or clinical populations may be limited. Second, although our use of cartoon representations substantially accelerates registration, it may not always achieve the accuracy of probabilistic approaches employed in the original Bayesian algorithm, e.g., in convoluted regions where the probabilistic labels are blurry and yield averaged intensities that may not faithfully represent the underlying tissue.

Overall, our new method offers a powerful and accessible tool for fast, high-resolution brain segmentation. By combining detailed histological priors with a lightweight implementation, it bridges the gap between expert-curated atlases and real-world neuroimaging needs. Its speed, simplicity, and robustness make it especially well-suited for large-scale studies at the subregion level. Crucially, by removing the need for specialized software or manual anatomical expertise, this freely available tool democratizes access to fine-grained brain segmentation, enabling a wider community of researchers to explore the brain at unprecedented spatial detail.

## Data and Code Availability

A ready-to-use tool is available as part of FreeSurfer: https://surfer.nmr.mgh.harvard.edu/fswiki/HistoAtlasSegmentation. As with the rest of FreeSurfer, the source code is available on GitHub: https://github.com/freesurfer/freesurfer/tree/dev/mri histo util.

## AuthorContributions

- Conceptualization: OP, YB, JEI
- Data curation: AC, MM, ER, JA, SC, EB, BB, AA, LZ, DLT, DK, MB
- Formal analysis: AC, LZ, OP, JEI
- Funding acquisition: OP, JEI
- Investigation: AC, MM, ER, LP, RA, JA, SC, EB, LZ, YB, OP, JEI
- Methodology: AC, MM, OP, YB, JLH, ZJ, JEI
- Project administration: ER, JLH, CS, ZJ, OP, JEI
- Resources: DLT, JLH, CS, ZJ
- Software: AC, MM, BB, AA, OP, YB, PS, JH, JN, RD, JEI
- Supervision: ER, LP, DK, MB, JLH, CS, ZJ, JEI
- Validation: AC, JEI
- Visualisation: OP, AC, PS, JH, JEI
- Writing – original draft: OP, JEI
- Writing – review and editing: all authors.

## Acknowledgements

The authors thank Rohit Jena for his prompt and helpful responses to our inquiries regarding his FireANTs method and software.

## Supplementary Material

### Heuristic intensity estimates for creating a synthetic MRI scan

We first compute the median intensities using the coarse segmentation from BrainFM for the white matter (*µ*^wm^), gray matter (*µ*^gm^), cerebellar white matter (*µ*^cwm^), cerebellar gray matter (*µ*^cgm^), caudate (*µ*^ca^), putamen (*µ*^pu^), and pallidum (*µ*^pa^). Next the average value and difference between white and gray matter medians is computed:

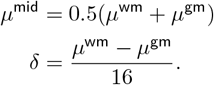

The substructure intensities where no direct label correspondence between the coarse BrainFM segmentation and NextBrain exists are created from *µ*^mid^ and *δ* as follows:

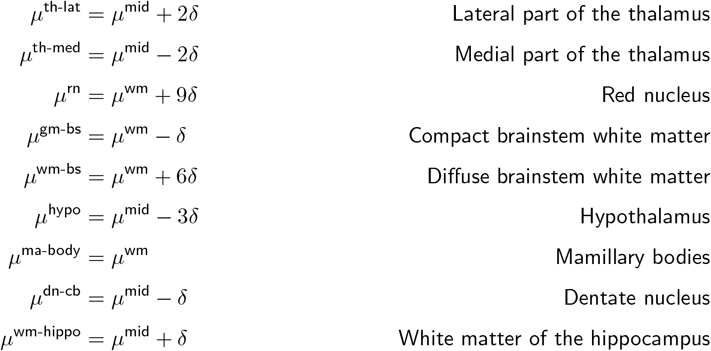

### Ex vivo comparison against the manual labeling on (Edlow et al., 2019)

**Table S1:**
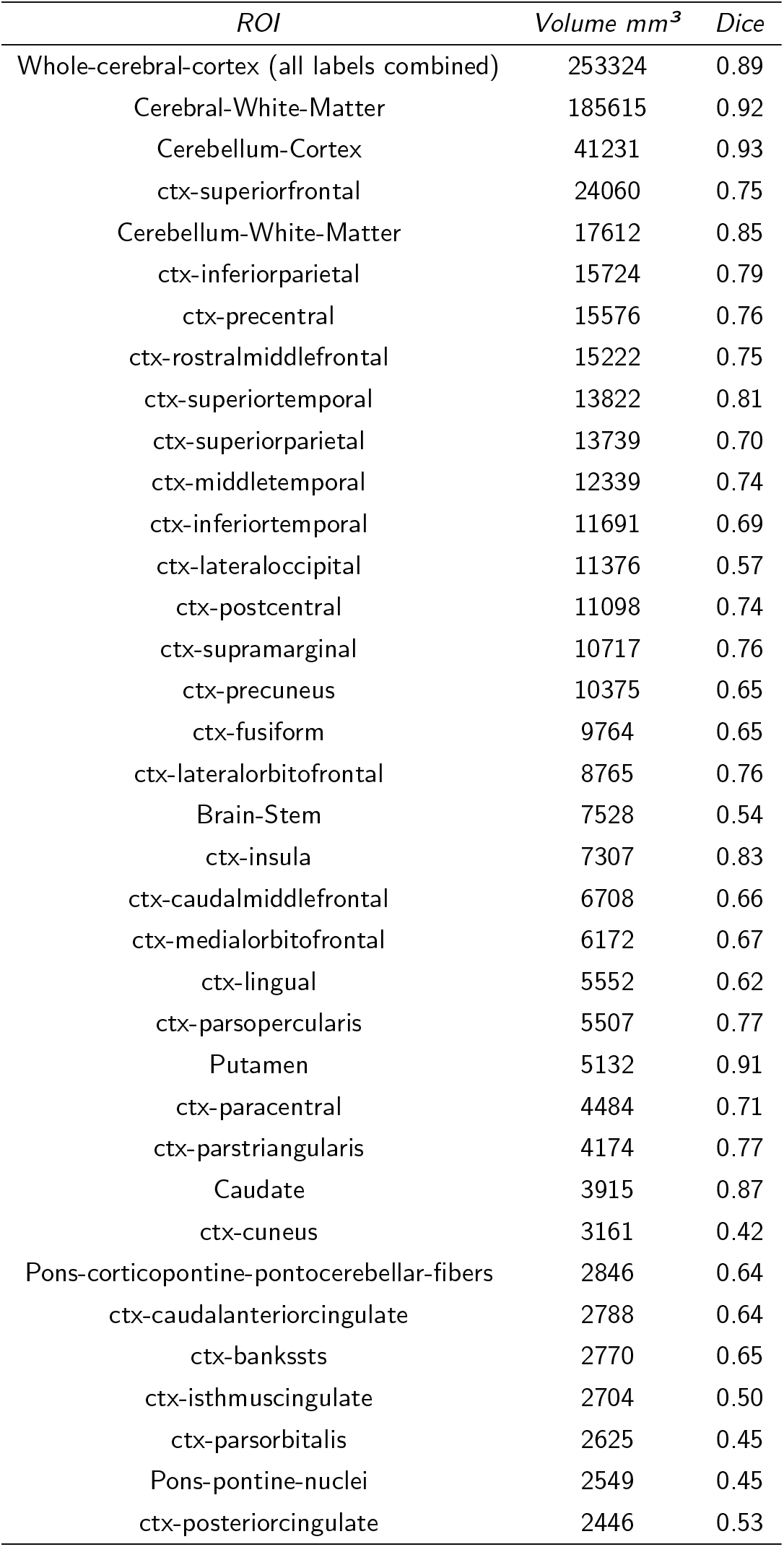
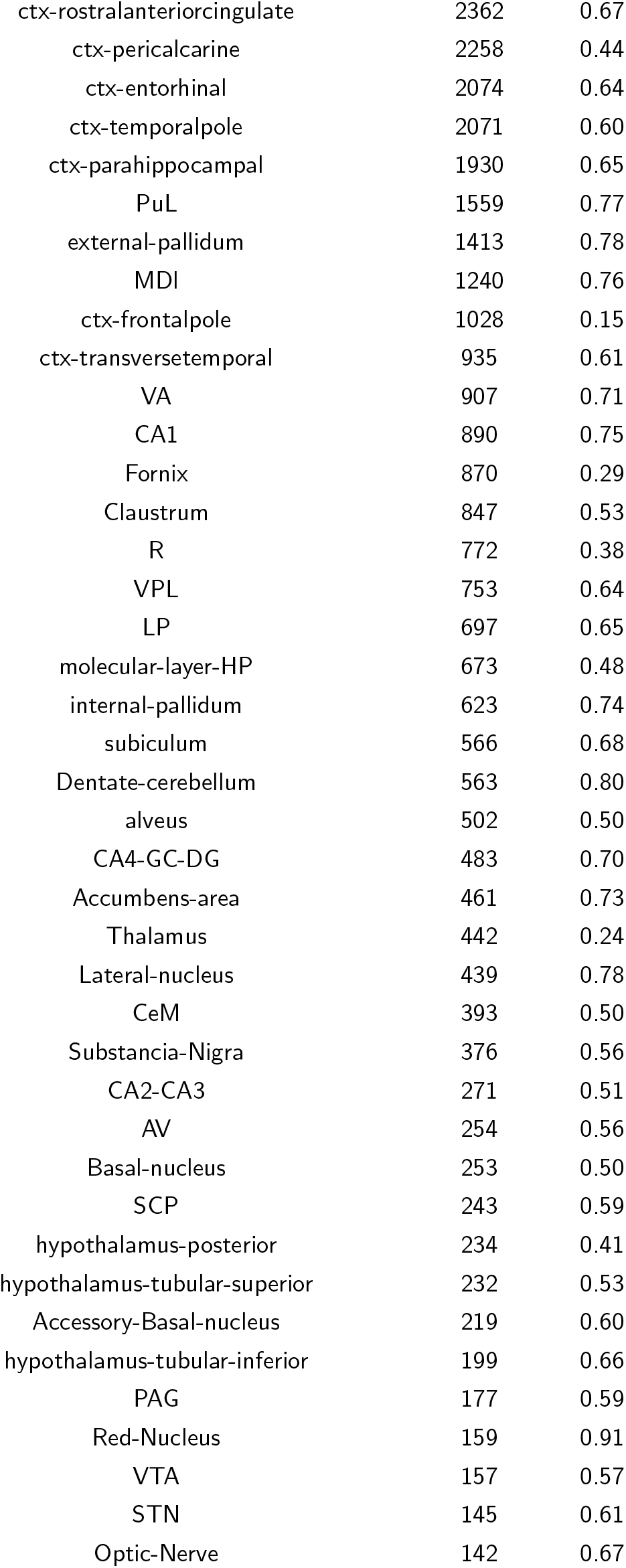
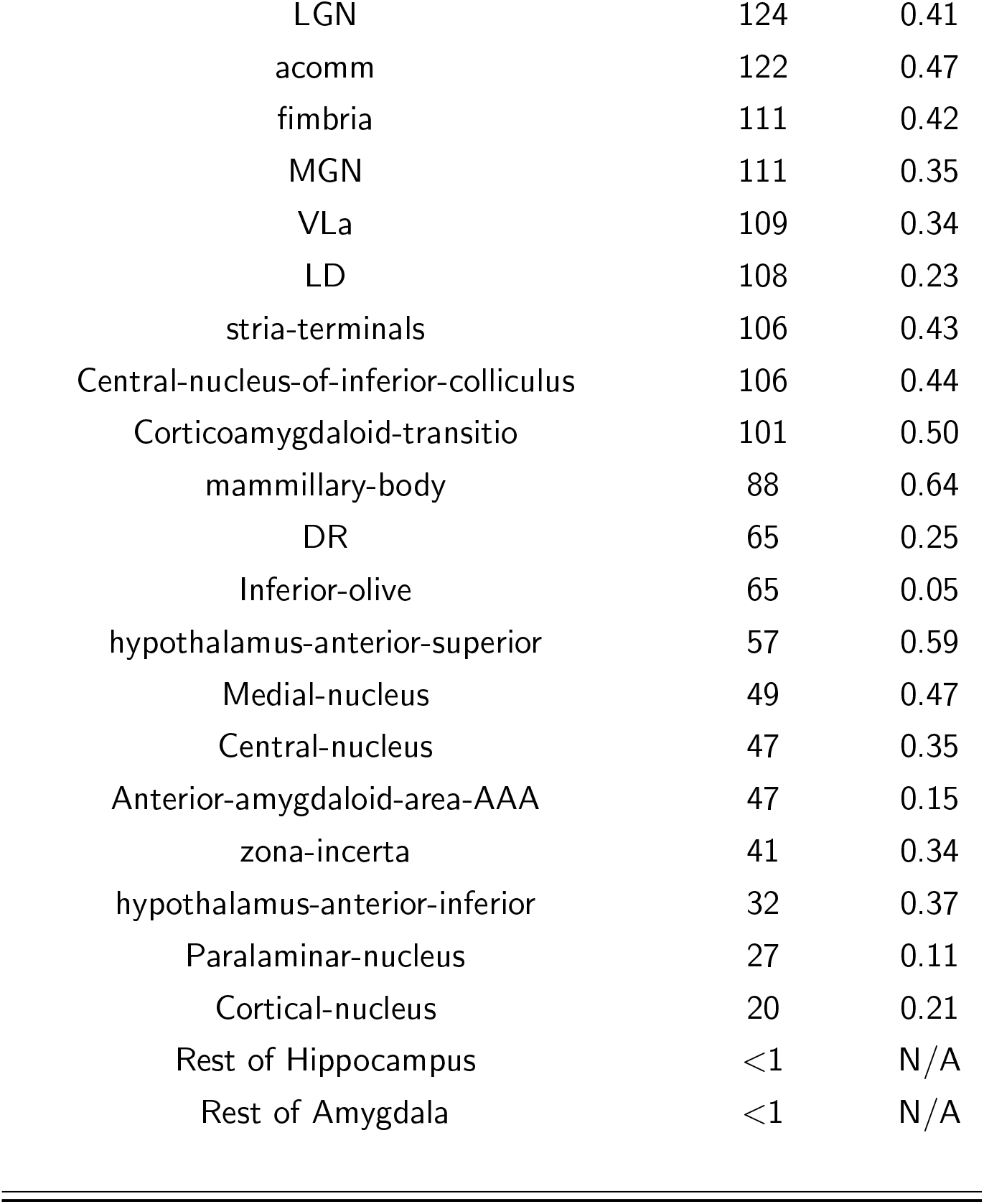
The volumes and Dice scores across all ROIs on the *ex vivo* sample from Edlow et al., 2019.

### Intra-class correlation analysis on the MIRIAD data set

**Table S2:**
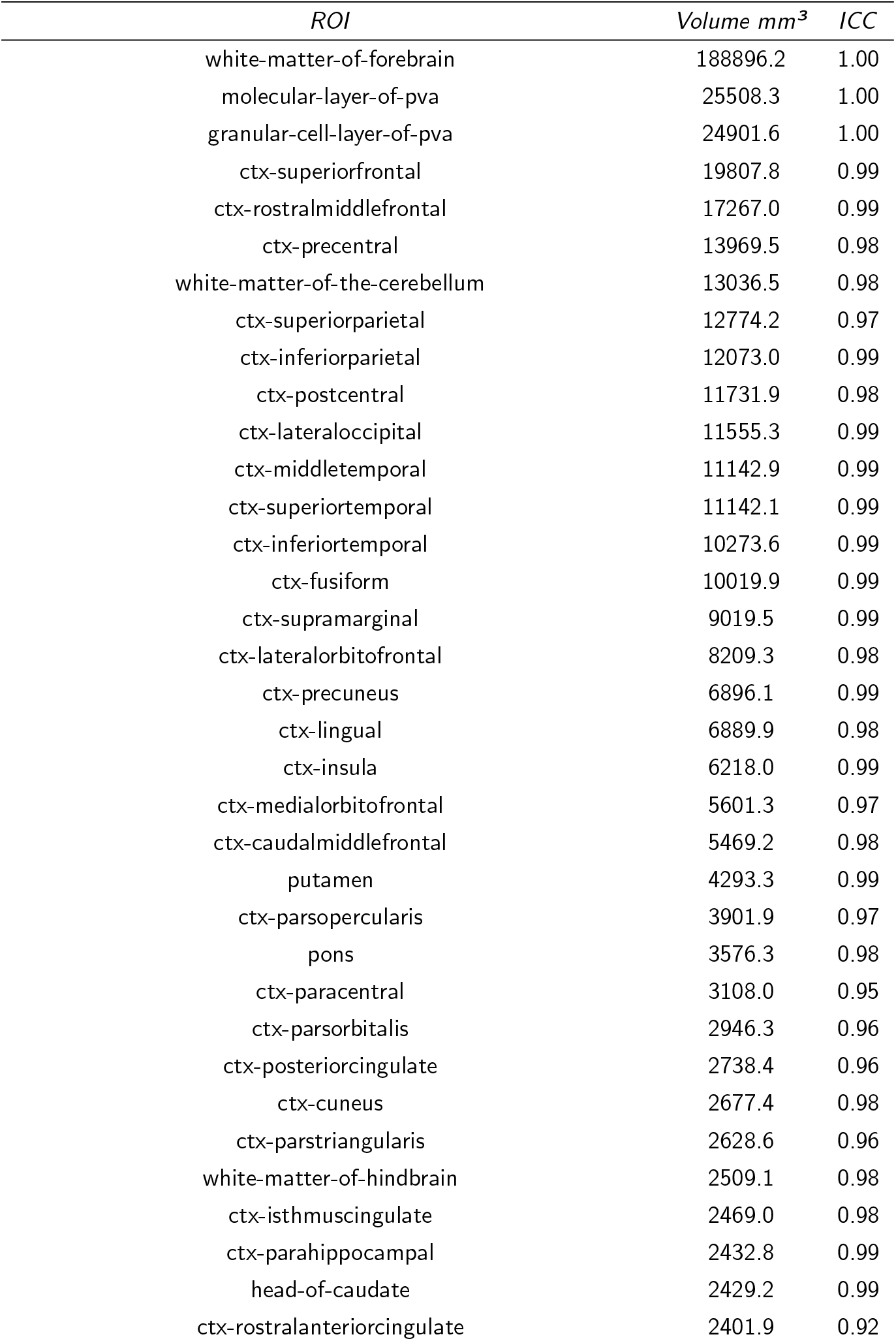
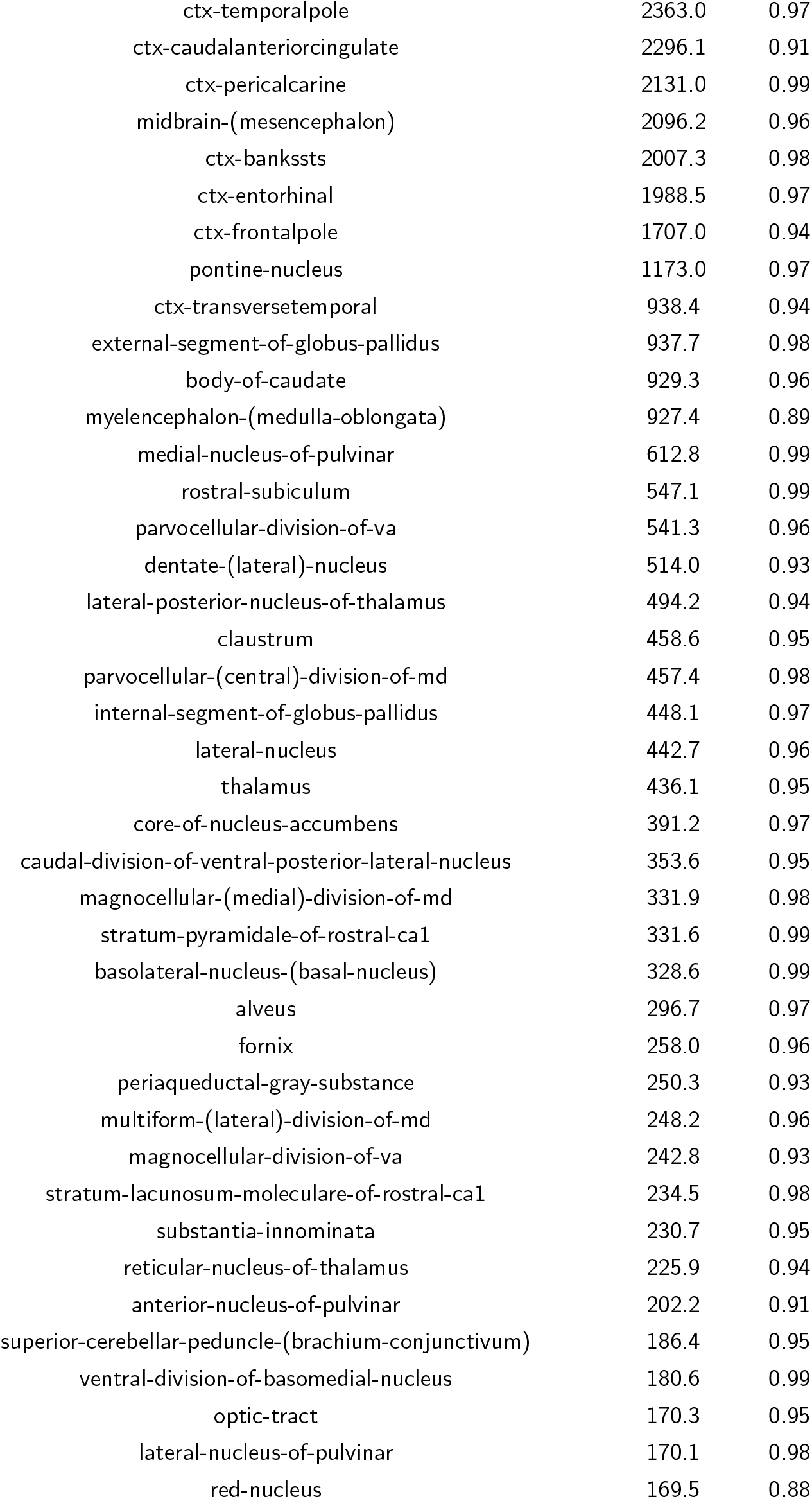
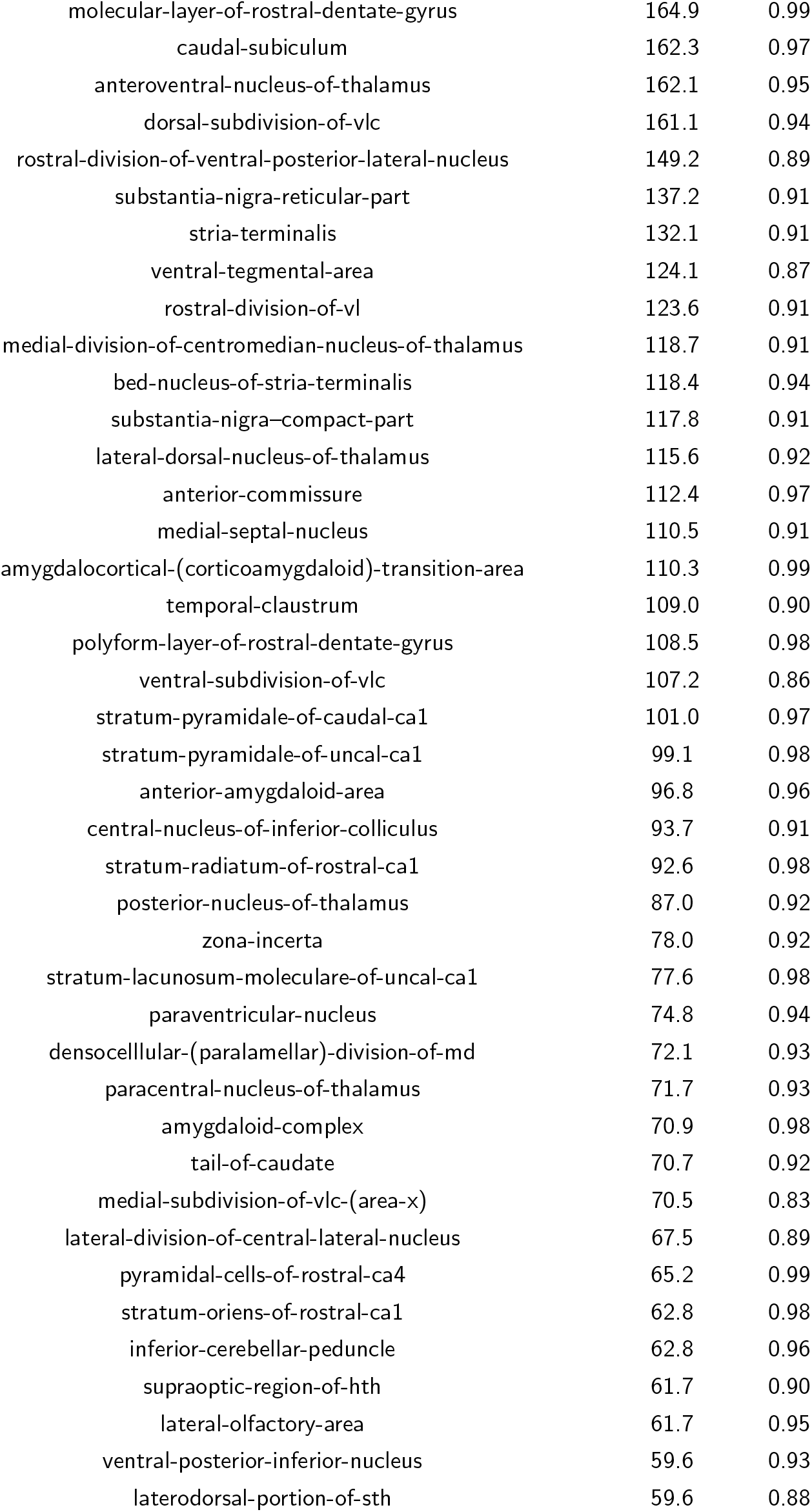
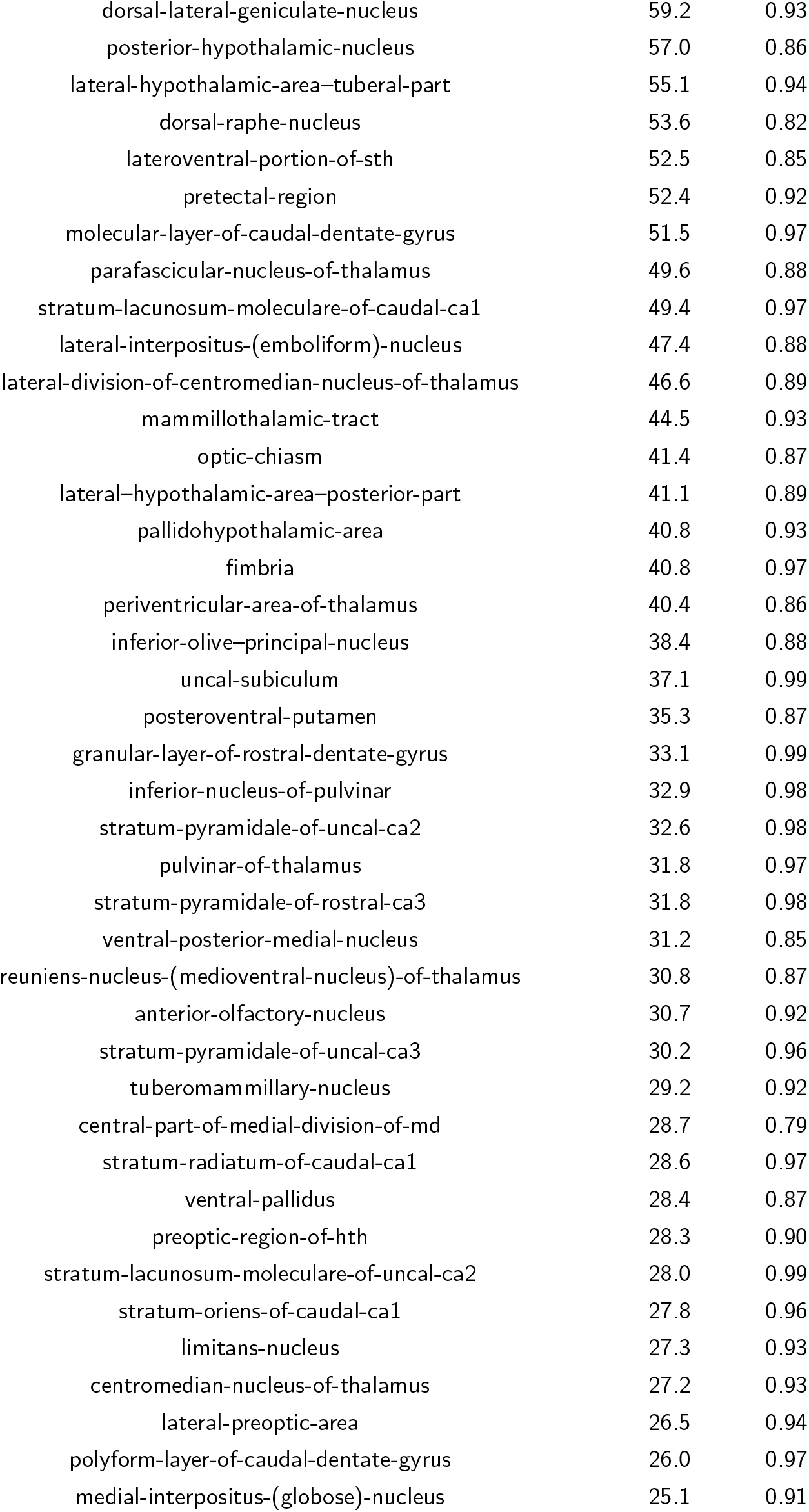
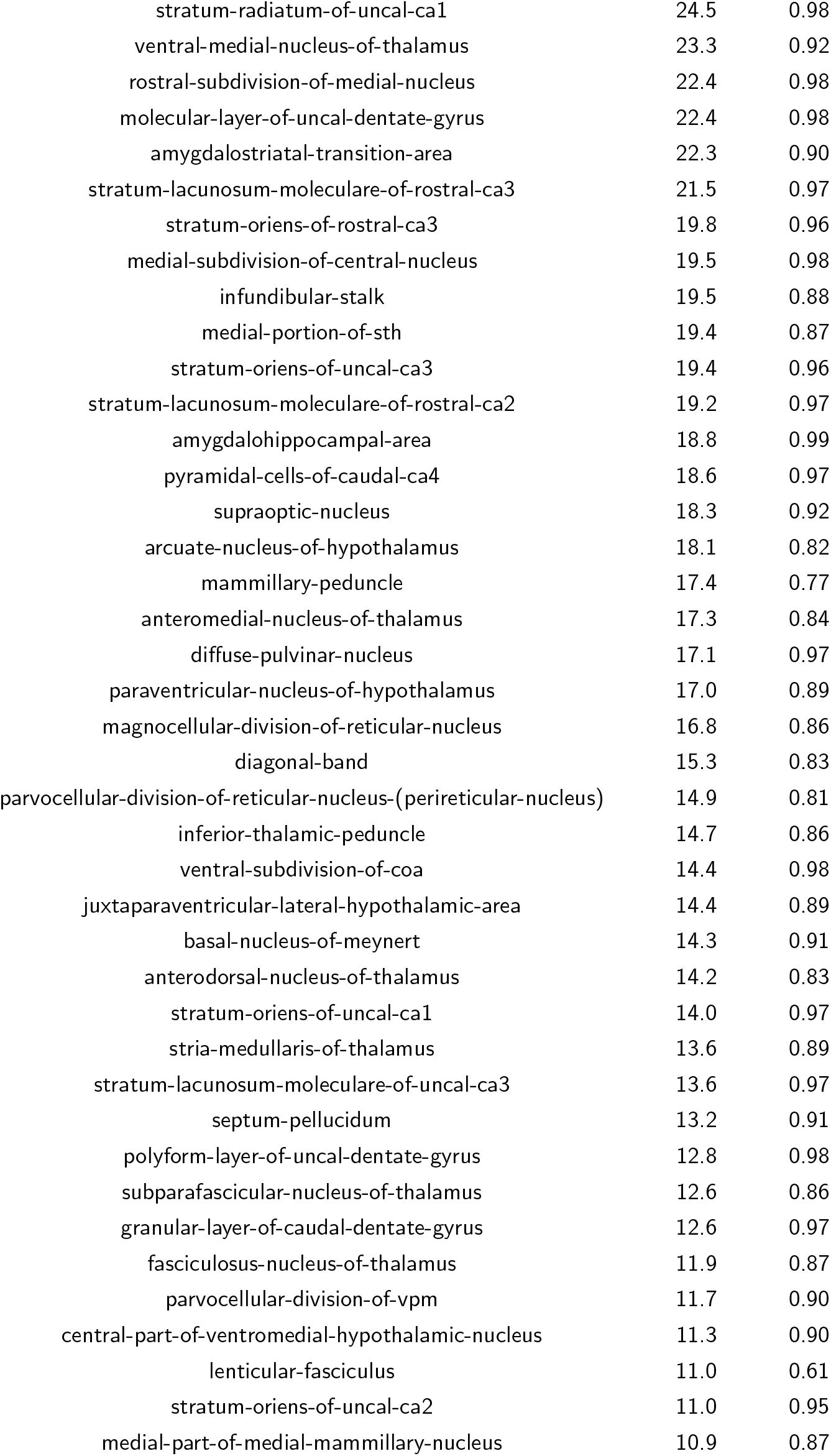
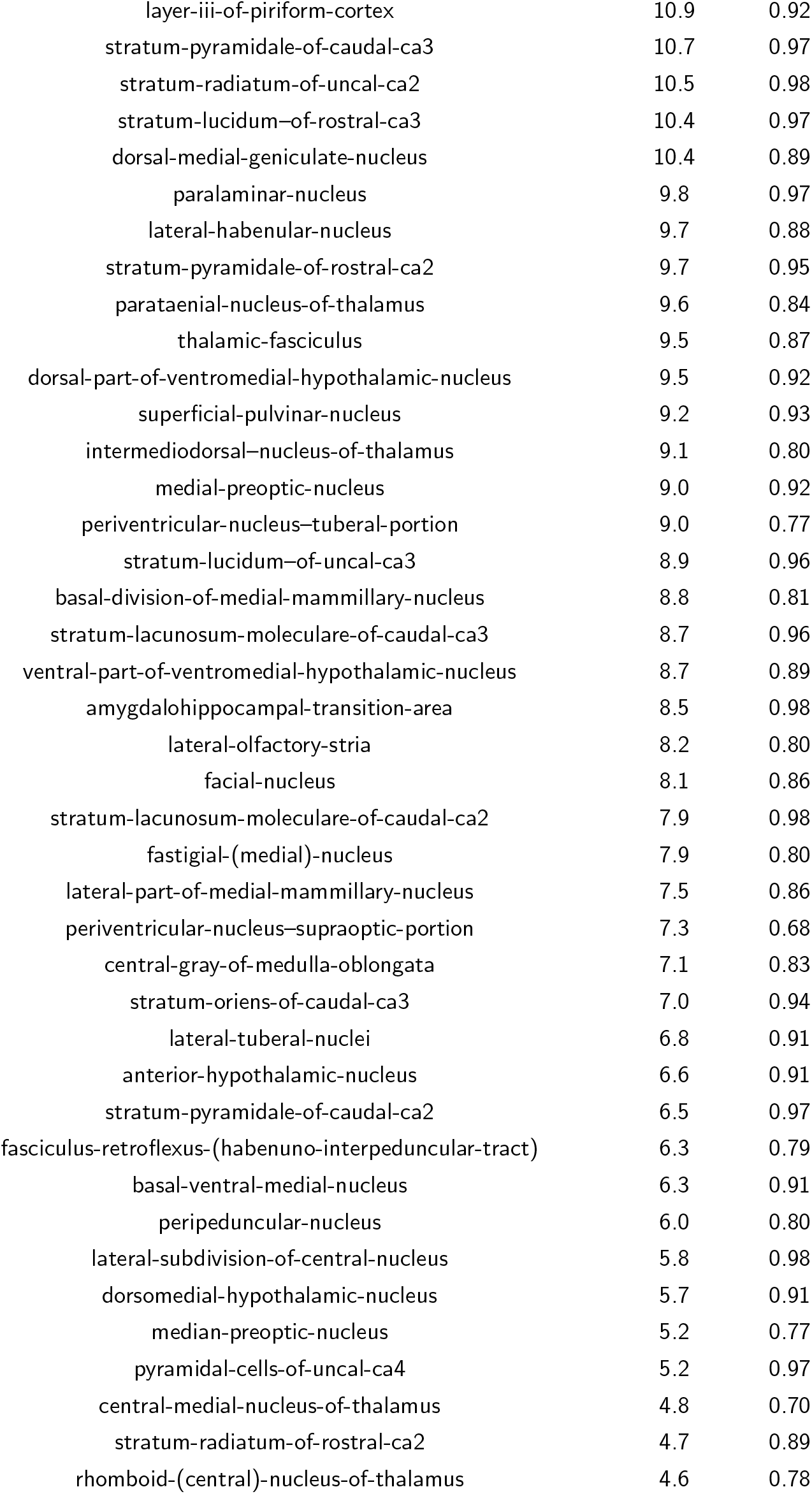
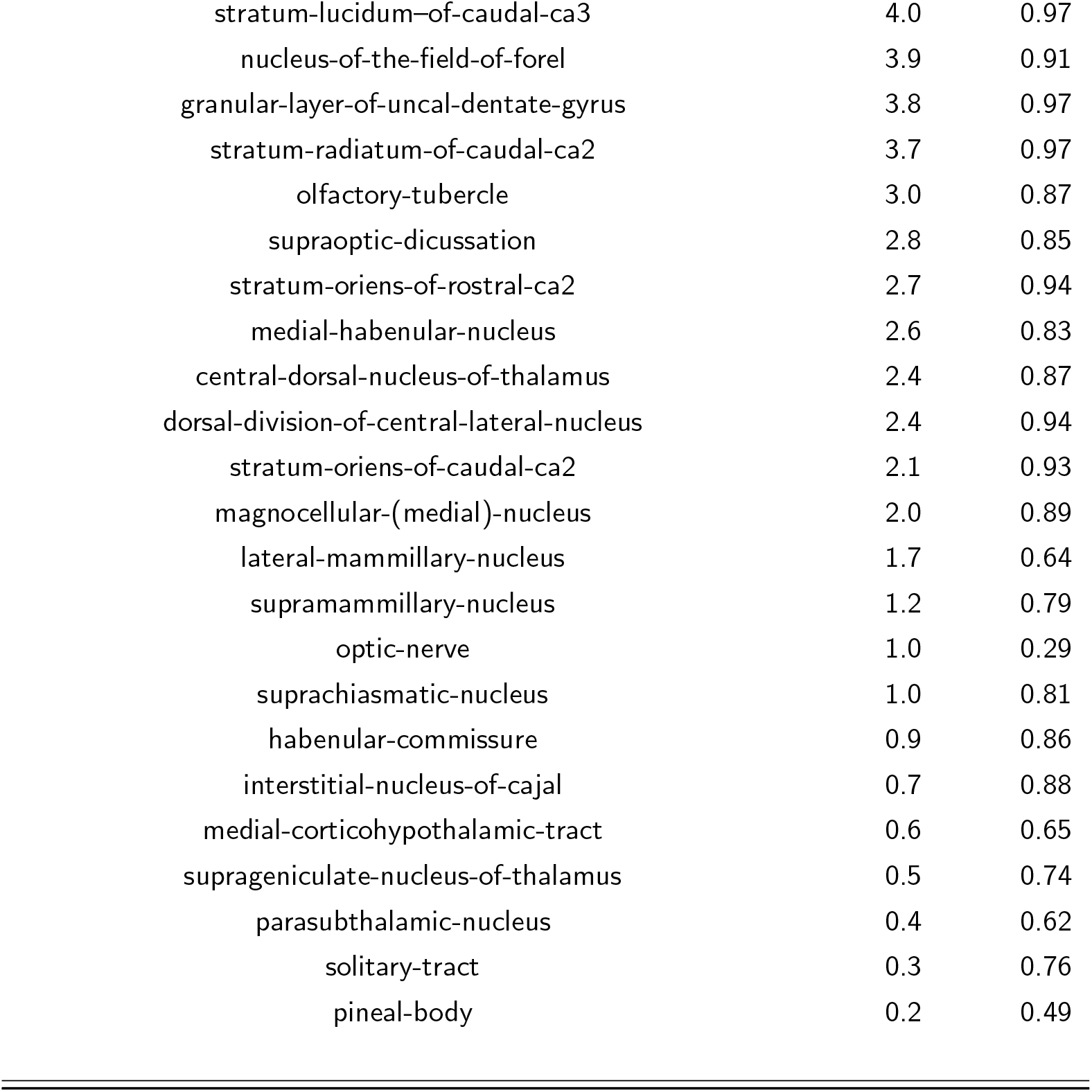
The volumes and intra-class correlation coefficients (ICC) over all the ROIs in the NextBrain atlas for the MIRIAD data set. The ROIs are listed in from largest to smallest.

